# Ligand-independent c-Met activation by HHLA2 drives hepatocellular carcinoma and predicts c-MET inhibitor efficacy

**DOI:** 10.1101/2024.11.07.622557

**Authors:** Xubo Huang, Runya Fang, Yuqian Pang, Zhe Zhang, Jieru Huang, Yingchang Li, Tao Yuan, Yuyi Zeng, Ziying Yao, Silvia Vega-Rubín-de-Celis, Josephine Thinwa, Hao Shen, Jiahong Wang, Feng Shen, Yongjie Wei

**Affiliations:** State Key Laboratory of Respiratory Disease, Affiliated Cancer Hospital & Institute, School of Basic Medical Science, Guangzhou Medical University, Guangzhou, China; Shenzhen Bay Laboratory & National-Regional Key Technology Engineering Laboratory for Medical Ultrasound, School of Biomedical Engineering, Shenzhen University Medical School, Shenzhen University; Clinical Research Institute and Department of Hepatic Surgery, Eastern Hepatobiliary Surgery Hospital, Shanghai, China; National Centre for Liver Cancer, Shanghai, China; Department of Hepatobiliary and Pancreatic Surgery, Tenth People’s Hospital of Tongji University, Shanghai, China; Institute of Cell Biology (Tumor Research), University Hospital Essen, Virchowstr. 173, 45122 Essen, Germany; Department of Internal Medicine, University of Texas Southwestern Medical Center, Dallas, Texas, USA

## Abstract

The HGF/c-Met signaling pathway facilitates the initiation, progression, and metastasis of hepatocellular carcinoma (HCC). c-Met activation, however, is complex and not solely dependent on HGF, hindering targeted therapy development. This study identifies a critical oncogenic role for HHLA2, a B7 family member, in HCC and highlights its potential as a therapeutic target. We demonstrate that HHLA2 directly interacts with and activates c-Met through N-glycosylation, triggering sustained signaling and promoting aggressive HCC features. Mechanistically, we identified the pro-tumorigenic role of HHLA2 required downstream upregulation of MMP9 and VEGFA, both implicated in tumor progression. In multiple mouse models, HHLA2 overexpression accelerated tumor progression, metastasis, and reduced liver NK cell infiltration, all of which were reversed by c-Met inhibition. In a cohort of 176 HCC patients, HHLA2 expression strongly correlated with c-Met phosphorylation, advanced tumor stage, and poor prognosis. Importantly, HHLA2 expression predicted sensitivity to c-Met inhibitors in cell lines and patient-derived organoids and could be detected in patient serum, suggesting its potential as a prognostic biomarker. Collectively, our findings reveal an HHLA2-mediated mechanism of c-Met activation and provide a strong rationale for targeting the HHLA2-c-Met axis as a novel therapeutic strategy, with HHLA2 serving as a potential prognostic biomarker.

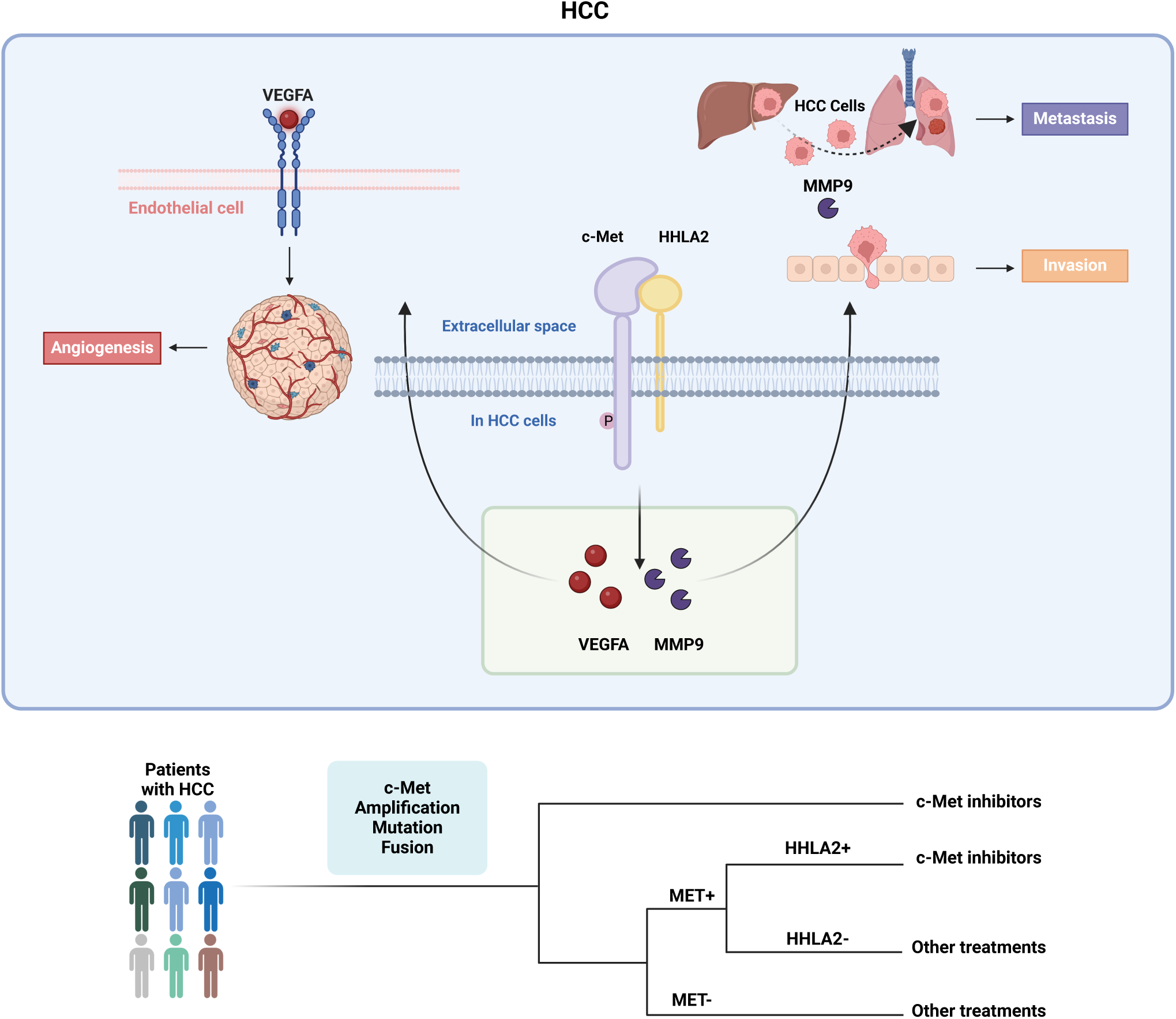

## Introduction

The HGF/c-Met signaling axis constitutes a critical regulatory hub governing hepatic physiology and pathobiology. Upon binding to its receptor tyrosine kinase c-Met, hepatocyte growth factor (HGF) initiates a complex signaling cascade indispensable for liver development, regeneration, and maintenance of homeostasis(1–4). Ligand engagement induces c-Met dimerization and subsequent autophosphorylation of specific tyrosine residues within its kinase domain (Tyr-1234 and Tyr-1235), thereby activating downstream signaling pathways, including Ras/MAPK, PI3K/Akt, and STAT(5, 6). A tightly regulated negative feedback loop involving c-Met ubiquitination and lysosomal degradation attenuates excessive signaling (7).

Dysregulation of the HGF/c-Met axis is implicated in HCC pathogenesis. Aberrant activation of this pathway stimulates cell proliferation, evades apoptosis, enhances cell motility, and induces angiogenesis, thereby promoting tumorigenesis. C-Met activation occurs through canonical HGF-mediated mechanisms and various atypical pathways, including receptor crosstalk, aberrant ligand engagement, and overexpression-induced autoactivation. These collective effects contribute to tumor initiation, growth, invasion, and metastasis, underscoring the pivotal role of c-Met in HCC progression (8–15).

While c-Met holds promise as a therapeutic target, developing effective therapies has been challenging and hindering clinical progress. This is exemplified by the disappointing clinical trial results of tivantinib, a highly anticipated c-Met inhibitor (9, 15–17). A key hurdle is identifying patients who would benefit, as c-Met inhibitors are generally effective only in tumors exhibiting "c-Met addiction," where cancer cells depend on c-Met activation for survival and growth (6). Currently, c-Met overexpression serves as a proxy for c-Met addiction in clinical trials due to the lack of effective biomarkers to directly measure c-Met activity in liquid biopsies. This, along with the complexity of c-Met activation, can lead to an inaccurate assessment of actual c-Met dependence(18, 19). To improve outcomes in HCC, we must fully understand the molecular mechanisms driving c-Met activation in this cancer. Critically, we must develop reliable, non-invasive biomarkers, significantly those detectable in liquid biopsies, to accurately predict patient response to c-Met targeted therapies.

Recent investigations have expanded the functional repertoire of the B7 family beyond its classical role in immune regulation, implicating these co-stimulatory molecules in the complex process of tumorigenesis(20, 21). Among the B7 family members, HHLA2 is a unique and intriguing player in tumorigenesis. Its evolutionary origin as a HERV-H endogenous retroviral protein, with a restricted expression profile to primates, underscores its distinct biological properties(22).

Initially identified for its expression on hematopoietic cells, HHLA2 has been increasingly recognized for its aberrant expression in various solid tumors, including HCC(22). While the exact mechanisms underlying the oncogenic potential of HHLA2 remain to be fully elucidated, accumulating evidence suggests its involvement in promoting tumor growth, invasion, and metastasis. These malignant properties may be attributed, in part, to HHLA2’s ability to interact with and modulate growth factor signaling pathways(23, 24).

Given the established role of the HGF/c-Met axis in HCC progression and the emerging oncogenic functions of HHLA2, exploring the potential interplay between these two pathways is warranted. Elucidating the molecular mechanisms underlying this interaction could reveal novel therapeutic targets and strategies for HCC.

This study elucidates a critical role for HHLA2 in driving HCC progression. Our findings demonstrate that HHLA2 directly interacts with c-Met, constitutively activating downstream signaling pathways and promoting a malignant phenotype characterized by increased proliferation, invasion, and angiogenesis. Notably, HHLA2 expression is closely correlated with advanced disease stages and poor patient prognosis, suggesting its potential as a prognostic biomarker. Targeting c-Met effectively inhibits the growth of HHLA2-positive HCC cells, highlighting a promising therapeutic strategy. These findings underscore the importance of HHLA2 as a novel therapeutic target for HCC.

## Results

### HHLA2 Expression Correlates with Aggressive Phenotype and Poor Prognosis in Hepatocellular Carcinoma

HHLA2, a B7 family member known for its immunosuppressive functions, exhibits minimal expression in most healthy tissues except the gastrointestinal tract (Supplemental Figure 1A). Conversely, HHLA2 is highly expressed in various malignancies, as evidenced by data from The Cancer Genome Atlas (TCGA) and the Protein Atlas database (Supplemental Figure 1B). Analysis of TCGA data revealed that HHLA2 mRNA levels were significantly elevated in tumor tissues, including HCC, compared to their corresponding normal tissues (Figure 1A). Furthermore, high HHLA2 expression in HCC patients significantly correlated with advanced tumor grade (Figure 1B) and worse overall survival (OS) (Figure 1C). Notably, multivariate Cox regression analysis confirmed HHLA2 as an independent prognostic factor for OS in this patient population (p=0.022) (Figure 1D).

**Figure 1.**
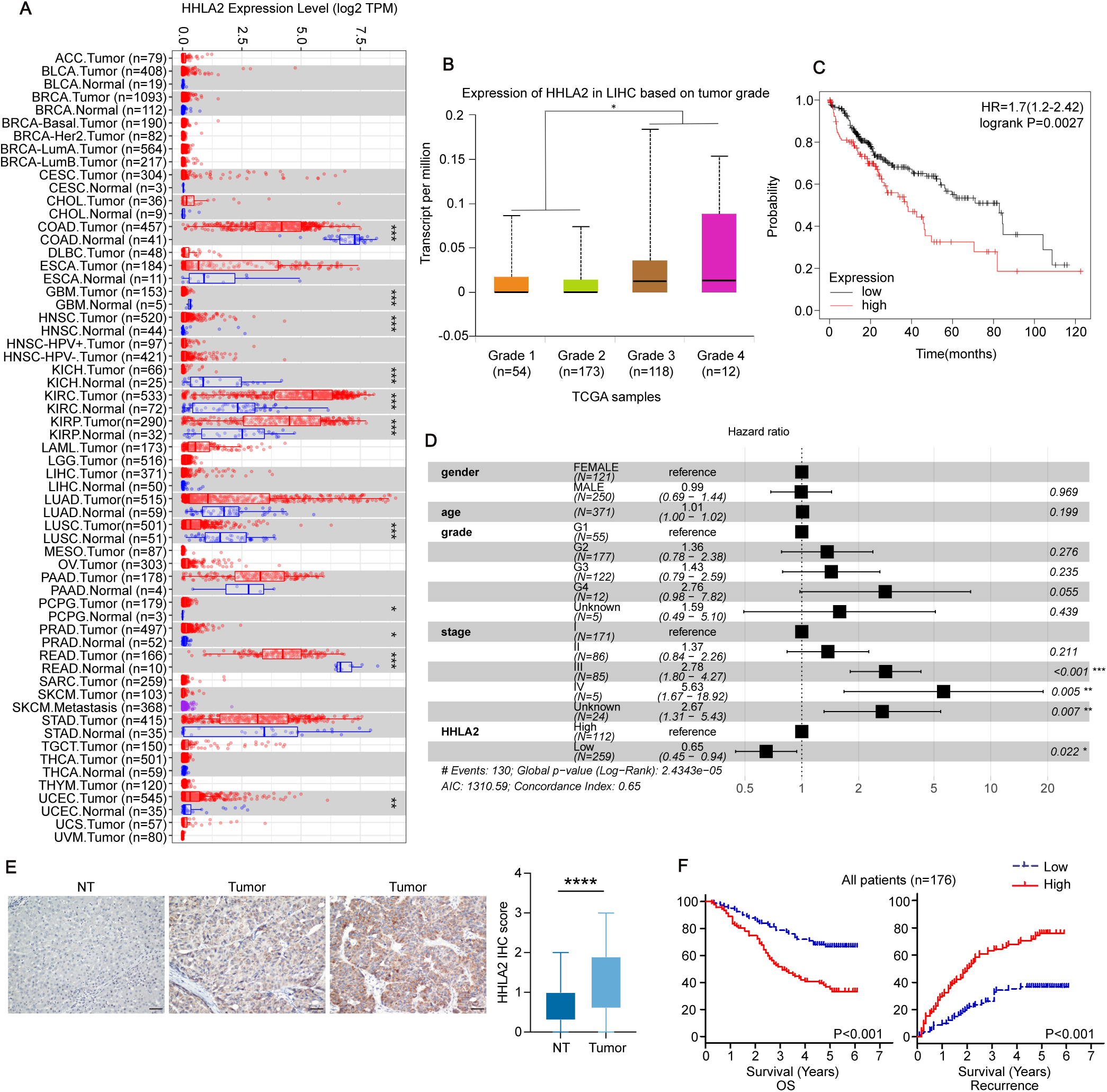
HHLA2 expression in HCC and its impact on patient prognosis. **(A, B)** HHLA2 mRNA expression in various tumor tissues (A) and in patients with different tumor grades of HCC (B) (n = 371 individuals). Data source: TCGA. TPM, transcripts per million. P values were determined by unpaired two-tailed t test. **(C, D)** Kaplan-Meier survival curves (C) and multivariate Cox regression analysis (D) of overall survival (OS) in patients with HCC (n = 359 individuals). Data source: TCGA. P values were determined by log-rank test (C) and Wald’s test (D). **(E)** HHLA2 protein levels in HCC and paired normal tissues assessed by immunohistochemical staining (n = 176 cases). P values were determined by paired two-tailed t test. **(F)** Kaplan-Meier survival curves of OS and recurrence in the HCC cohort (n = 176 individuals). P values were determined by log-rank test. * P < 0.05, ** P < 0.01, *** P < 0.001, **** P < 0.0001.

To validate these observations and elucidate the clinical significance of HHLA2 expression in HCC, we performed immunohistochemical analysis on tumor tissues from a cohort of 176 HCC patients in Guangdong, China. Our results corroborated the TCGA data, demonstrating significantly higher HHLA2 protein levels in tumor tissues compared to adjacent non-tumor tissues (Figure 1E). Patients with high HHLA2 expression exhibited a more aggressive clinical profile, characterized by larger tumors (p < 0.001), increased vascular invasion (p = 0.016), advanced Barcelona Clinic Liver Cancer (BCLC) stages (p < 0.001), and higher recurrence and metastasis rates (p < 0.001) (Supplemental Figure 1C, Supplemental Table 1). No significant associations were found with gender, age, alpha-fetoprotein (AFP) levels, hepatitis B surface antigen (HBsAg) status, cirrhosis, tumor number, satellite nodules, tumor capsule status, or tumor differentiation (Supplemental Table1). Importantly, survival analysis (Figure 1F, Supplemental Table 2) and both univariate and multivariate Cox regression analyses identified HHLA2 as an independent risk factor for poor prognosis in HCC patients (Hazard Ratio [HR] = 2.758; 95% Confidence Interval [CI] = 1.719-4.426) (Supplemental Table 1, 2)

To elucidate the functional role of HHLA2 in HCC progression, we employed a panel of HCC cell lines with varying endogenous HHLA2 expression levels. HepG2 and Hep3B cells displayed low HHLA2 mRNA and protein levels, while Huh7 cells exhibited comparatively high HHLA2 expression (Supplemental Figure 2A, 2B). We generated stable cell lines with either ectopic HHLA2 expression (HepG2-HHLA2 and Hep3B-HHLA2) or HHLA2 deficiency (Huh7-HHLA2KD) using shRNA (Supplemental Figure 2C, 2D).

Overexpression of HHLA2 significantly promoted proliferation in HepG2 and Hep3B cells by day 5, as measured by the CCK8 assay (Figure 2A, Supplemental Figure 2F). Conversely, HHLA2 knockdown inhibited Huh7 cell proliferation. This effect was reversed upon reintroduction of HHLA2 (Supplemental Figure 2G, 3A). Similarly, HepG2-HHLA2 and Hep3B-HHLA2 cells displayed enhanced colony formation (Supplemental Figure 2H, 2I), increased anchorage-independent growth in soft agar assays (Supplemental Figure 2K), faster migration (Figure 2B, Supplemental Figure 3B), and greater invasion capabilities in transwell assays (Figure 2C, Supplemental Figure 3D) compared to control cells. These pro-tumorigenic abilities were significantly reduced in Huh7-HHLA2KD cells but restored upon HHLA2 re-expression (Supplemental Figure 2J, 3C, 3E, 3G).

**Figure 2.**
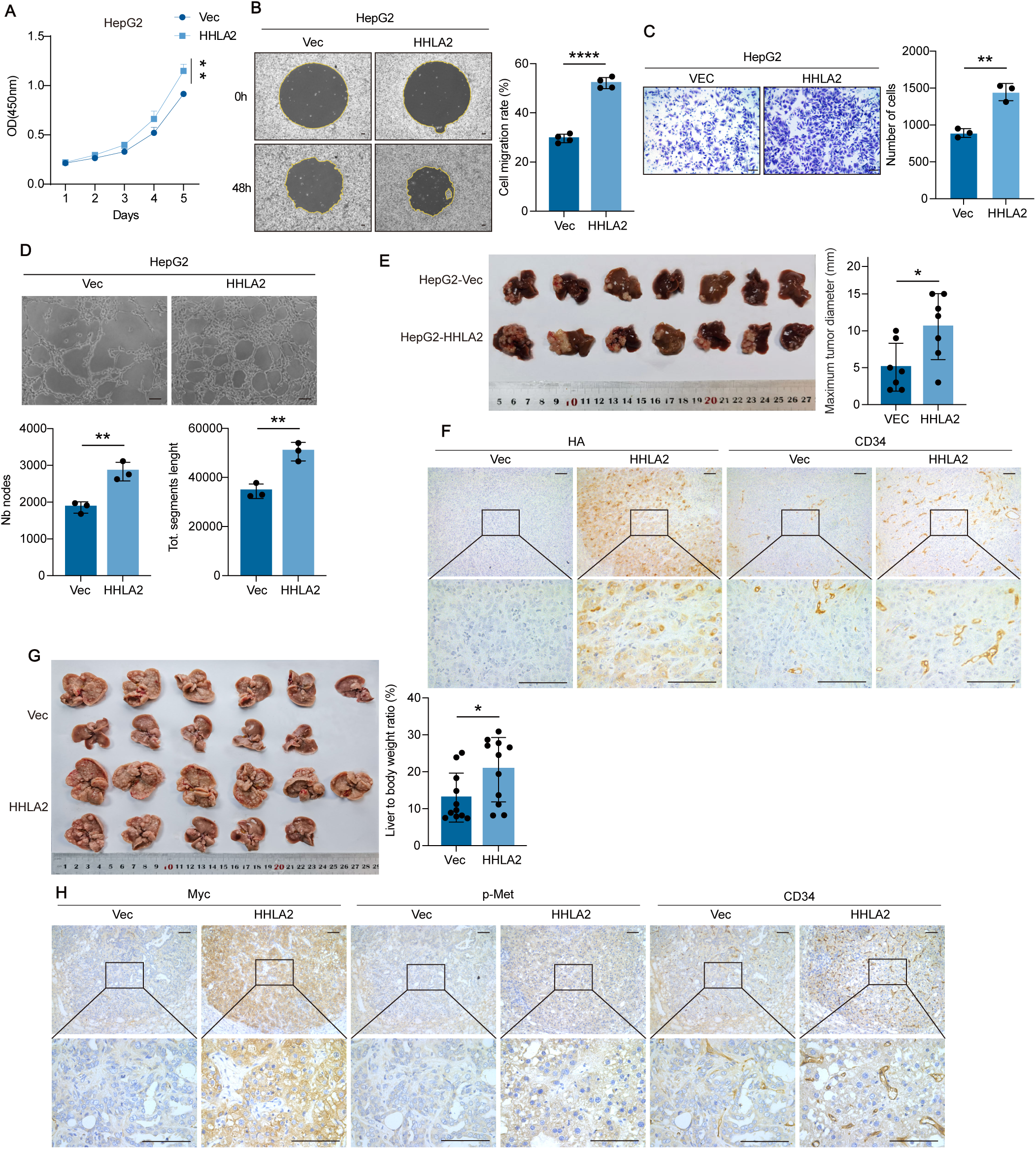
HHLA2 promotes aggressive phenotypes in HCC. **(A–D)** Assays for tumorigenic phenotypes of HepG2-Vec and HepG2-HHLA2 cells: (A) cell proliferation assessed by CCK-8 assay; (B) cell migration assay; (C) Transwell invasion assay; (D) tube formation assay in HUVECs using conditioned media (CM) from HepG2 cells. **(E, F)** Representative images (E) and immunohistochemical staining for CD34 (blood vessels) and HA (HHLA2) (F) of liver tumors in orthotopic xenograft models in nude mice injected with HepG2-Vec or HepG2-HHLA2 cells (n = 7 mice per group). **(G, H)** Representative images and quantification (liver/body weight ratio) (G) and immunohistochemical staining for CD34 (blood vessels), pY1235-Met (activated c-Met), and myc (HHLA2) (H) of orthotopic liver tumors in C57BL/6 mice following hydrodynamic tail vein injection (HDTVi) of myc-HHLA2 or a control vector, along with N-RasV12/myr-AKT1 and Sleeping Beauty transposon system (n = 11 mice per group). P values were determined by one-way ANOVA (A) or two-tailed Student’s t test (B–H). * P < 0.05, ** P < 0.01, *** P < 0.001, **** P < 0.0001. Scale bars, 100 μm.

HCC development relies on tumor angiogenesis, a critical process supplying oxygen and nutrients for tumor progression (25). To investigate HHLA2’s role in this process, we cultured human umbilical vein endothelial cells (HUVECs) with conditioned media (CM) from HCC cells and assessed their capillary-like tube formation(26). Compared to the control group, HUVECs treated with CM from HepG2-HHLA2 and Hep3B-HHLA2 cells displayed a significant increase in total tube length and number of branching nodes (Figure 2D, Supplemental Figure 3F). Conversely, HHLA2 knockout in Huh7 cells markedly reduced the capacity of their CM to promote HUVEC tube formation. This effect was reversed entirely upon HHLA2 reintroduction (Supplemental Figure 3G).

Next, we explored the role of HHLA2 in HCC progression in vivo using two mouse liver cancer models. In the orthotopic liver xenograft model, nude mice injected with HepG2-HHLA2 cells exhibited increased tumor diameters compared to controls (Figure 2E). Conversely, mice injected with Hu7H-HHLA2KD cells displayed the opposite phenotype (Supplemental Figure 3H). For the second model, we employed hydrodynamic tail vein injection (HDTVi) to deliver HA-HHLA2 or a control vector, along with activated N-RasV12 and myr-AKT1 (myristylated AKT1) and Sleeping Beauty transposase into the livers of immunocompetent C57BL/6 mice, thereby inducing tumor formation(27). By week 4.5, both HHLA2 and control mice developed multiple tumor lesions. However, the liver-to-body weight ratio indicated a significantly higher tumor burden in the HHLA2-expressing mice (Figure 2G). Immunohistochemical staining for CD34, a marker of vascular endothelial progenitor cells, on liver tissues from both models revealed higher microvessel density in tissues expressing HHLA2, confirming its pro-angiogenic role in HCC (Figure 2F, 2H).

Furthermore, we replaced N-RasV12/myr-AKT1 with c-Met or c-Met/myr-AKT1 in the HDTVi model to generate HCC with different genetic backgrounds. While tumor development took longer in these mice (around 12 weeks for c-Met and 8 weeks for c-Met/myr-AKT1), HHLA2 expression significantly promoted HCC formation in both cases (Supplemental Figure 3I, 3J).

### HHLA2 Interacts with and Constitutively Activates c-Met

To investigate the mechanism by which HHLA2 promotes HCC progression, we analyzed RNA-sequencing (RNA-seq) data from TCGA and protein expression data from The Cancer Proteome Atlas (TCPA). Elevated HHLA2 expression was found to be correlated with increased levels of p-Met, p-MEK1, eIF4E, and c-Myc, all indicative of enhanced downstream signaling. Concurrently, a decrease in the levels of p-AMPK and p-GSK3, known tumor suppressors in HCC, was also observed (Supplemental Figure 4A). Interestingly, the Kyoto Encyclopedia of Genes and Genomes (KEGG) pathway analysis revealed that differentially expressed genes between HHLA2 high and HHLA2 low HCC were significantly enriched in the PI3K-AKT and MAPK signaling pathways (Supplemental Figure 4B).

Next, we performed RNA-seq transcriptome analysis in HepG2 cells. Overexpression of HHLA2 significantly altered the transcription of 796 genes, with 244 genes upregulated and 552 genes downregulated (|log2(fold change) | > 0.4, p < 0.05) (Supplemental Figure 4C). Similar to the TCGA and TCPA analyses, KEGG enrichment analysis again demonstrated significant enrichment of genes influenced by HHLA2 expression in the PI3K-AKT and MAPK pathways (Figure 3A). Furthermore, Gene Set Enrichment Analysis (GSEA) indicated that these differentially expressed genes are enriched in processes promoting cell proliferation, such as DNA endoreduplication and positive regulation of spindle checkpoint (Supplemental Figure 4D). These findings suggest that HHLA2 may promote HCC progression by enhancing these signaling pathways.

**Figure 3.**
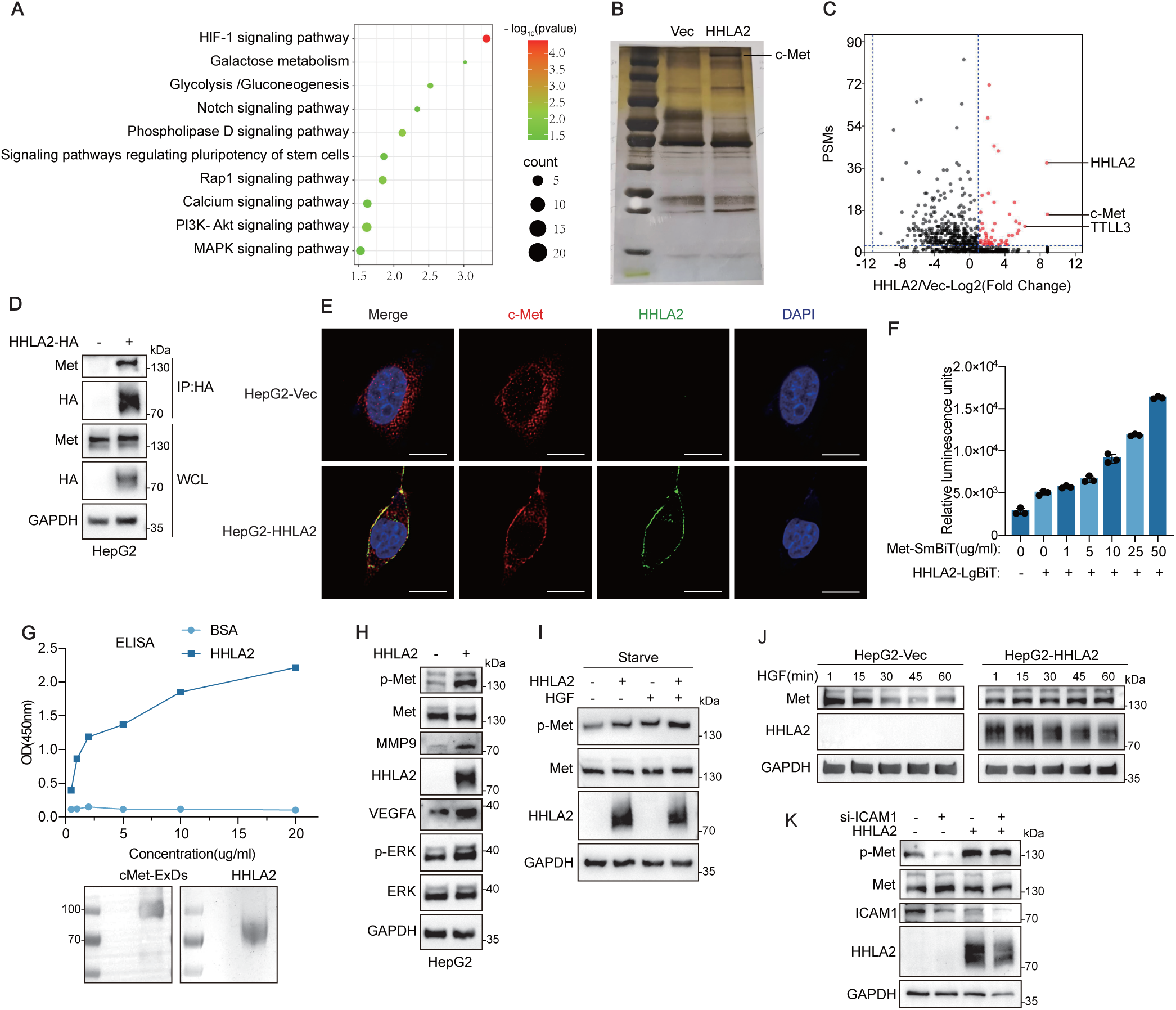
HHLA2 interacts with and constitutively activates c-Met. **(A)** KEGG pathway enrichment analysis of the HepG2 transcriptome after HHLA2 overexpression. **(B)** Silver-stained SDS-PAGE gel of anti-HA immunoprecipitates from HepG2-HHLA2-HA and HepG2-Vec cells. **(C)** Volcano plot depicting the differential abundance of proteins identified by mass spectrometry in HHLA2 versus vector immunoprecipitates from (B). The fold change in protein abundance (HHLA2/vector) is plotted on the x axis against peptide-spectrum match [PSM]) on the y axis. **(D)** Coimmunoprecipitation of HHLA2-HA and c-Met with anti-HA antibody in HepG2 cells with or without HHLA2 overexpression. **(E)** Representative immunofluorescence images of HHLA2 and c-Met in HepG2-Vec (upper panel) and HepG2-HHLA2 (lower panel) cells. Cells were serum starved overnight and then stimulated with 1 μM HGF for 15 minutes before fixation. Scale bar, 4 μm. **(F)** Split luciferase complementation assay to define the interaction domains between HHLA2 and c-Met. Luminescence was measured following the addition of indicated amounts of recombinant c-MET-ExD-SmBit-6His protein to 293-LgBiT-HHLA2 cells. **(G)** In vitro binding assay to examine the HHLA2 and c-Met interaction. Indicated amounts of HHLA2 ExD-SmBit-6His recombinant protein or BSA control were added to 96-well plates precoated with 1 μ g/well of purified c-MET-ExD-SmBit-6His protein and incubated for 1 hour. Following washes, the amount of HHLA2 protein bound to c-Met was detected by ELISA. The lower panel shows the input purified c-MET-ExD-SmBit-6His and HHLA2 ExD-SmBit-6His recombinant proteins. **(H, I)** Western blot analysis of the indicated proteins in HepG2 cells with or without HHLA2 overexpression under normal conditions (H) or when cells were serum starved and then stimulated with 40 ng/ml HGF for 30 minutes (I). **(J)** Analysis of c-Met degradation following activation: HepG2-Vec or HepG2-HHLA2 cells were serum starved overnight and then stimulated with 40 ng/ml HGF for the indicated times. **(K)** Western blot analysis of the indicated proteins in HepG2-Vec or HepG2-HHLA2 cells following ICAM1 knockdown.

To identify potential proteins interacting with HHLA2 in HCC, we immunoprecipitated HHLA2 from HepG2-HHLA2 cells and performed label-free, semi-quantitative mass spectrometry analysis. This analysis revealed significant changes in the abundance of co-immunoprecipitated proteins upon HHLA2 overexpression (Figure 3B, 3C). Notably, c-Met and STT3 emerged as prominent candidate interactors. C-Met signaling is well-established to play a pivotal role in various cellular processes, including proliferation, survival, and migration. Upon binding its ligand, hepatocyte growth factor (HGF), c-Met undergoes phosphorylation of crucial tyrosine residues within its cytoplasmic domain, triggering downstream PI3K/AKT and MAPK signaling pathways(5). Our identification of HHLA2 interacting with c-Met aligns with our previous transcriptomic analyses, suggesting that HHLA2 promotes these pathways.

To validate a potential physical interaction between HHLA2 and c-Met, we employed co-immunoprecipitation assays in HepG2 and Hep3B cells. The results showed a reciprocal interaction, with HHLA2 pulling down c-Met and vice versa (Figure 3D, Supplemental Figure 5A, 5B). This interaction was further supported by confocal microscopy, which revealed co-localization of c-Met and overexpressed HHLA2 at the cell membrane in HepG2 cells (Figure 3E).

We then employed a split-luciferase complementation assay(28) to define the domains involved in the interaction. HEK293 cells expressing LgBiT-tagged c-Met demonstrated a dose-dependent increase in luminescent signal upon the addition of recombinant SmBiT-tagged HHLA2-ExD (extracellular domain), indicating direct binding between their extracellular regions. Similar results were obtained when HEK293 cells expressing LgBiT-tagged HHLA2 were incubated with c-Met-ExD harboring a SmBiT tag (Figure 3F, Supplemental Figure 5C). Next, we employed complementary approaches to validate this interaction in a cell-free system. Utilizing a secretory expression system in suspension-cultured 293f cells, we independently expressed and purified the ExDs of HHLA2 and c-Met. Subsequently, an ELISA assay utilizing c-MET-coated plates confirmed the direct interaction between HHLA2 and c-MET (Figure 3G). Notably, the ELISA results demonstrated a dose-dependent increase in HHLA2 binding to c-Met with increasing protein concentration. These findings provide strong evidence for a direct interaction between HHLA2 and c-MET mediated through their extracellular domains.

We investigated the potential role of HHLA2 in c-MET activation. HHLA2 may function in two ways: as a co-receptor, facilitating HGF binding to c-Met, or as an agonist, directly activating the receptor. The short cytoplasmic domain of HHLA2 suggests a more likely role as an agonist, with c-Met’s cytoplasmic domain responsible for recruiting downstream signaling molecules. Supporting this hypothesis, overexpression of HHLA2 in HepG2 and Hep3B cells significantly increased c-Met phosphorylation at Y1235 (a key activation site). This phosphorylation event was accompanied by activation of the downstream effector Erk and upregulation of MMP9 and VEGFA, both implicated in tumor progression (Figure 3H, Supplemental Figure 5G).

Furthermore, under serum starvation conditions that deplete HGF and other c-MET ligands, c-Met phosphorylation was abolished entirely in control HepG2-Vec cells. In contrast, c-Met in HepG2-HHLA2 cells remained phosphorylated at Y1235, suggesting ligand-independent activation by HHLA2 (Figure 3I). These findings were further validated in Huh7 cells. HHLA2 knockdown significantly reduced phosphorylated c-Met, phosphorylated ERK, MMP9, and VEGFA levels in normally cultured Huh7 cells. However, these effects were reversed upon HHLA2 replenishment (Supplemental Figure 5H). Moreover, Huh7 cells with replenished HHLA2 exhibited notable c-Met phosphorylation even under serum-starved conditions (Supplemental Figure 5I). These results collectively suggest that HHLA2 can directly activate c-Met in a ligand-independent manner.

We investigated their respective effects on receptor endocytosis and degradation to further elucidate the mechanisms underlying HHLA2-mediated c-Met activation compared to HGF stimulation. HGF binding typically triggers c-Met internalization and subsequent lysosomal degradation, which maintains signaling homeostasis(7). Consistent with these reports, serum-starved HepG2 cells treated with HGF exhibited c-Met internalization (Figure 3E), accompanied by a gradual decrease in total c-Met protein levels over time, as evidenced by Western blot analysis (Figure 3J). In contrast, HepG2 cells overexpressing HHLA2 displayed enhanced c-Met membrane localization (Figure 3F) and resistance to degradation, maintaining stable c-Met protein levels (Figure 3J). These findings suggest that HHLA2-mediated c-Met activation stabilizes the receptor at the plasma membrane, thereby preventing its endocytosis and subsequent lysosomal degradation. Consequently, this promotes sustained c-Met signaling, unlike the transient activation observed with HGF treatment alone.

Previous studies have shown that c-Met activation in HepG2 cells depends on its co-receptor, ICAM-1(29). We investigated whether HHLA2 could functionally substitute for ICAM1 in this role. In HepG2-Vec cells with ICAM1 knockdown, HGF-induced c-Met phosphorylation was abolished, indicating the requirement of ICAM-1 for ligand-dependent c-Met activation (Figure 3K). In contrast, HepG2-HHLA2 cells exhibited sustained c-Met phosphorylation even under serum-starved conditions, where the availability of the HGF ligand would be significantly reduced (Figure 3K). These findings indicate that HHLA2 can functionally compensate for the absence of ICAM-1, facilitating ligand-independent c-Met activation.

### N-Glycosylation of HHLA2 Maintains Its Interaction with c-Met

HHLA2 is known to be heavily glycosylated by the STT3 oligosaccharyltransferase complex in colorectal cancer, impacting its stability, cell surface localization, and immune evasion(30). Glycosylation patterns can influence protein function, and HHLA2 in HepG2 cells exhibits a characteristic smeared band pattern on western blots, indicative of extensive glycosylation and potentially higher molecular weight (Figure 3H, 3I). This finding, coupled with identifying STT3A as an HHLA2-interacting protein (Figure 3C), led us to hypothesize that glycosylation might regulate HHLA2 function in HCC, potentially affecting c-Met binding and activation. To examine this hypothesis, we treated HepG2-HHLA2 cells with Tunicamycin, an inhibitor of N-linked glycosylation. This treatment effectively abolished HHLA2 glycosylation, as demonstrated by a shift in its electrophoretic mobility on western blots (Supplemental Figure 5D) and a concomitant reduction in c-MET binding. We employed a complementary approach to confirm the observed effects were not due to potential Tunicamycin-mediated de-glycosylation of c-Met. We purified un-glycosylated HHLA2-SmBiT recombinant protein from Tunicamycin-treated 293f cells (Supplemental Figure 5F) and assessed its ability to interact with c-Met in a split-luciferase complementation assay. Incubation of HEK293 cells expressing LgBiT-tagged c-Met with un-glycosylated HHLA2-SmBiT did not induce luminescence emission. In contrast, glycosylated HHLA2-SmBiT from untreated cells readily did so (Supplemental Figure 5E). These findings demonstrate that glycosylation of HHLA2 is essential for its interaction with c-Met.

### HHLA2/c-Met Signaling Promotes HCC Progression via MMP9 and VEGFA

c-Met activation primarily engages the PI3K-AKT and RAS-MAPK signaling pathways, with ERK phosphorylation within the latter serving as a critical driver of tumorigenesis(29). Consistent with this, HHLA2 overexpression augmented ERK phosphorylation in HepG2 and Hep3B cells (Figure 3H, Supplemental Figure 5G). In contrast, its knockdown attenuated ERK activation in Huh7 cells, an effect rescued by HHLA2 re-expression (Supplemental Figure 5H, I). To determine the requirement for functional c-Met in HHLA2-mediated ERK activation, we employed the non-tumorigenic LO2 hepatocyte cell line, which expresses deficient levels of endogenous c-Met. Overexpression of c-MET in LO2 cells induced its autophosphorylation and activated its downstream signaling molecule, ERK. Interestingly, the co-expression of c-Met with HHLA2 further enhanced the phosphorylation of both c-Met and ERK. Importantly, expression of the kinase-deficient c-Met KD mutant (Y1234,1235F) alone or co-expression with HHLA2 did not induce ERK phosphorylation in LO2 cells (Figure 4A). These findings, along with the observation that c-Met KD abrogates the HHLA2-driven tumor-promoting effects in HDTVi HCC mouse models (Figure 5G) strongly support the notion that HHLA2-dependent ERK activation requires functional c-MET and its kinase activity.

**Figure 4.**
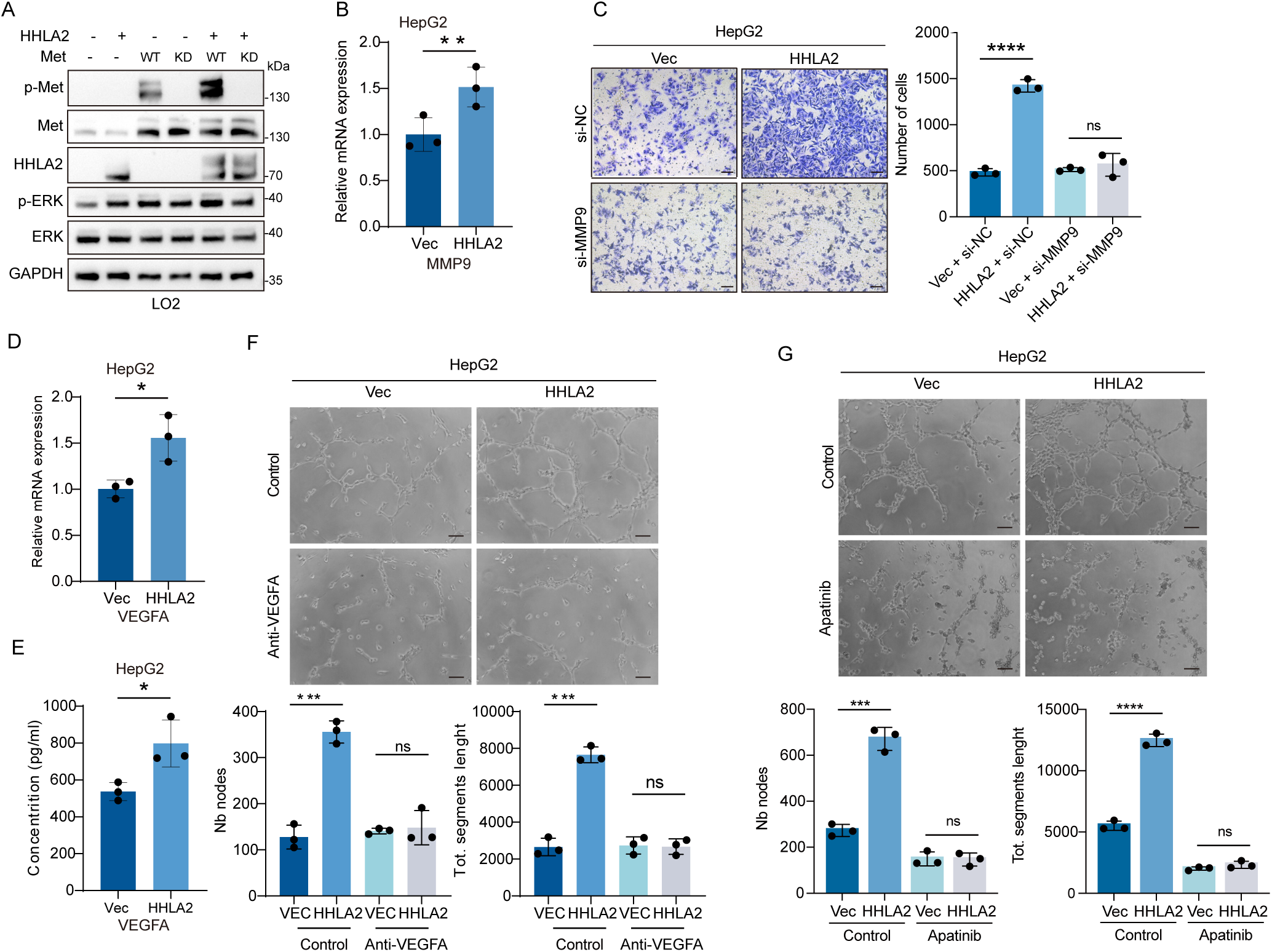
MMP9 and VEGFA mediate HHLA2-promoted invasion and tumor angiogenesis, respectively. **(A)** Western blot analysis of the indicated proteins in LO2 cells cotransfected with HHLA2 and c-MET (wild-type or kinase-dead) in the combinations shown. **(B)** qRT-PCR analysis of MMP9 mRNA expression in HepG2 cells with or without HHLA2 expression. **(C)** Transwell invasion assays of HepG2-Vec and HepG2-HHLA2 cells with or without MMP9 knockdown. **(D, E)** VEGFA mRNA expression assessed by qRT-PCR (D) and VEGFA secretion assessed by ELISA (E) in HepG2-Vec and HepG2-HHLA2 cells. **(F, G)** HUVEC tube formation assays using conditioned media from HepG2-Vec or HepG2-HHLA2 cells, with or without VEGFA depletion using an anti-VEGFA antibody (F) or in the presence or absence of 50 ng/ml apatinib (VEGF receptor inhibitor) (G). P values were determined by two-tailed Student’s t test. * P < 0.05, ** P < 0.01, *** P < 0.001, **** P < 0.0001. Scale bars, 100 μm.

**Figure 5.**
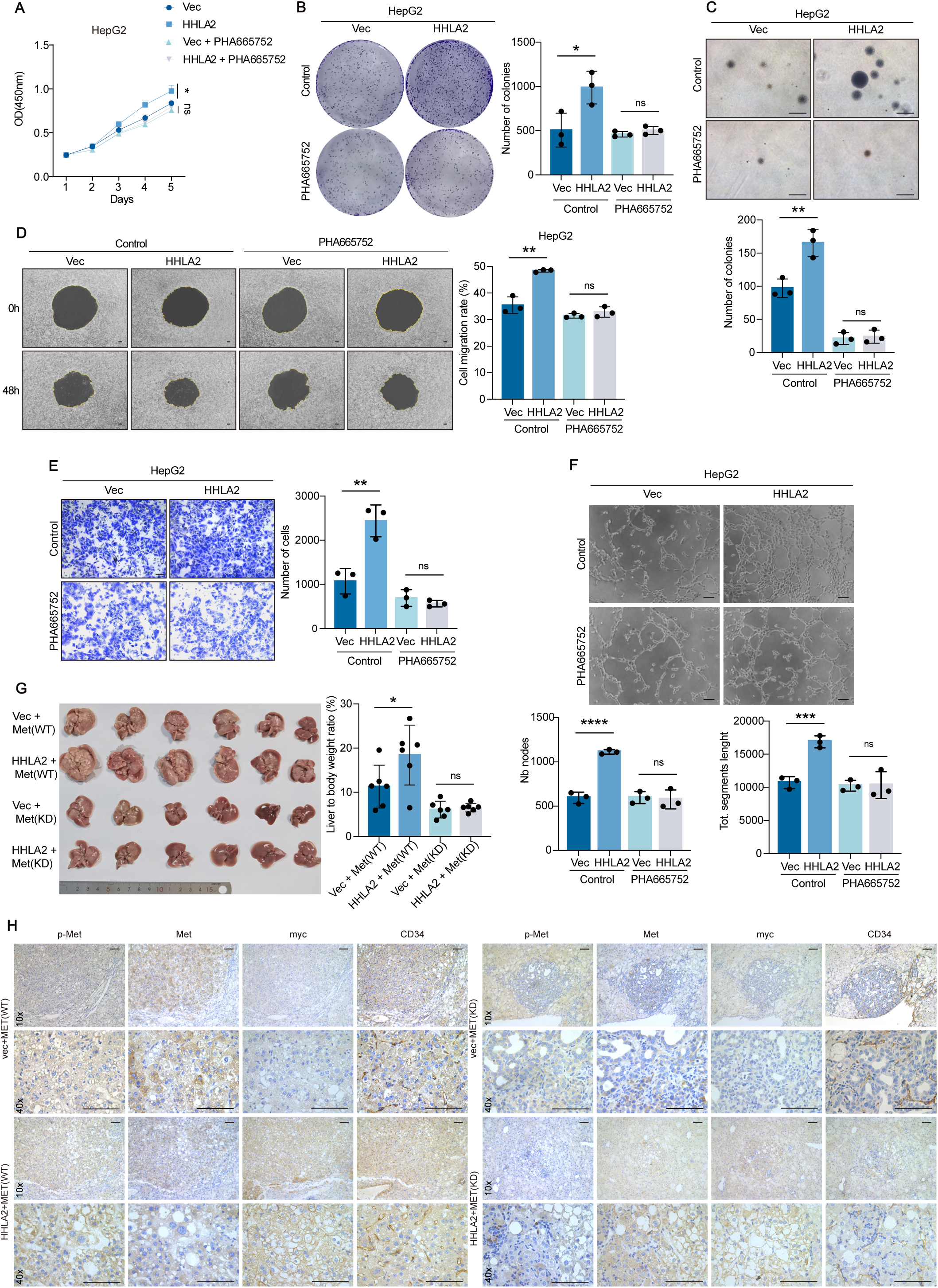
c-Met activation is indispensable for HHLA2-promoted HCC progression. **(A–F)** Assessment of malignant phenotypes in HepG2-Vec and HepG2-HHLA2 cells, with or without 100 nM c-Met inhibitor PHA665752 treatment. Phenotypes evaluated include (A) cell proliferation by CCK-8 assay, (B) two-dimensional colony formation, (C) soft agar colony formation, (D) cell migration, (E) Transwell invasion, and (F) HUVEC tube formation with conditioned media. (G) Representative images of orthotopic liver tumors and quantification of tumor burden (liver/body weight ratio) in C57BL/6 mice following HDTVi of HHLA2-myc or control vector, along with N-RasV12/myr-AKT1, c-MET (wild-type or kinase-dead), and Sleeping Beauty transposon system (n = 6 mice per group). **(H)** Immunohistochemical staining of tumor tissues from G for phosphorylated c-Met (p-Met; pY1235, indicative of activated c-Met), total c-Met, myc (HHLA2 marker), and CD34 (blood vessel marker). P values were determined by one-way ANOVA (A) or two-tailed Student’s t test (B–G). * P < 0.05, ** P < 0.01, *** P < 0.001, **** P < 0.0001. Scale bars, 100 μm.

Given that the most prominent phenotypes of HHLA2 in promoting HCC progression are metastasis and angiogenesis, we subsequently investigated its involvement in critical metastatic processes: epithelial-mesenchymal transition (EMT) and matrix metalloproteinase (MMP)-mediated extracellular matrix degradation, both known to be regulated by the PI3K-AKT and RAS-MAPK signaling pathways. Overexpression of HHLA2 in HepG2 cells did not induce EMT, as evidenced by unaltered E-cadherin and N-cadherin expression and the absence of morphological changes (Supplemental Figure 6A-C). While HHLA2 did not affect MMP2 and MMP7 expression (Supplemental Figure 6D), it significantly upregulated MMP9 expression (Figure 4B, Supplemental Figure 6H). Conversely, HHLA2 knockdown suppressed MMP9 expression in Huh7 cells, an effect rescued by HHLA2 re-expression (Supplemental Figure 6I), suggesting a causal link between HHLA2 and MMP9. To investigate the functional significance of MMP9 in HHLA2-mediated invasion, we employed siRNA-mediated MMP9 knockdown in HepG2 cells (Figure 4C, Supplemental Figure 6J). This effectively abolished the invasion-promoting effects of HHLA2 overexpression (Figure 4C), whereas MMP2 knockdown had no impact (Supplemental Figure 6E-G). Similarly, MMP9 knockdown reduced the invasiveness of Huh7 cells but did not further impair invasion in Huh7-HHLA2KD cells (Supplemental Figure 6K), confirming the role of MMP9 as a key downstream effector of HHLA2-driven invasion.

VEGF-A, a key tumor-derived angiogenesis factor, and its receptor VEGFR constitute the most prominent signaling pathway driving tumor angiogenesis. The enrichment of VEGF signaling genes, such as VEGFA, PTGS2, and PLCG2, among the upregulated genes in HepG2 cells, (Supplemental Figure 4C) upon HHLA2 overexpression further suggests the pivotal role of VEGFA in HHLA2-mediated HCC angiogenesis. In line with these findings, stable overexpression of HHLA2 in HepG2 and Hep3B cells triggered a substantial increase in VEGFA mRNA levels, protein abundance, and secretion (Figure 4D, 4E, 3H, Supplemental Figure 6L, 6N, 5G). Conversely, HHLA2 knockdown in Huh7 cells markedly decreased VEGFA parameters, restored upon HHLA2 re-expression (Supplemental Figure 5H, 6M, 6O). In vitro tube formation assays demonstrated that depletion of VEGFA from HepG2-conditioned media using a specific antibody significantly attenuated the pro-angiogenic effects of HHLA2 overexpression (Figure 4F). Similarly, inhibition of VEGFR2 signaling with Apatinib abolished the enhanced tube formation induced by HHLA2 (Figure 4G, Supplemental Figure 6P). These findings collectively indicate that HHLA2 promotes tumor angiogenesis by upregulating VEGFA expression and secretion from HCC cells, activating VEGFR2 signaling in endothelial cells.

### C-Met Activation Is Indispensable for HHLA2-Promoted HCC Progression

To elucidate whether HHLA2-mediated pro-HCC effects are contingent upon c-Met activation, we impaired c-Met activity using pharmacological or genetic approaches. We examined the impact of HHLA2 expression on tumor progression. c-Met inhibitor PHA665752 effectively blocked HHLA2 expression-induced ERK activation, VEGFA, and MMP9 upregulation in HepG2, Hep3B and Huh7 cells (Supplemental Figure 7A-D, F-H). Furthermore, the HHLA2-mediated pro-tumorigenic phenotypes in these HCC cells, including enhanced proliferation, 2-D and 3-D colony formation, migration, invasion, and angiogenesis, were significantly suppressed by PHA665752 treatment (Figure 5A-F, Supplemental Figure 8A-I). siRNA-mediated c-Met knockdown recapitulated the inhibitory effects of PHA665752 on HHLA2-driven HCC progression (Supplemental Figure 7E, 7I, 7J, Supplemental Figure 9A-F).

To validate the *in vivo* requirement for c-Met in HHLA2-mediated tumorigenesis, we employed the HDTVi mouse model of HCC. Mice harboring N-RasV12 and either wild-type (WT) or kinase-dead (KD) c-MET were co-injected with either HHLA2 or a control vector (VEC). Compared to the RasV12+c-Met+VEC group, mice injected with RasV12+ c-MET+HHLA2 developed significantly more aggressive liver tumors. Notably, replacing WT c-MET with the KD mutant completely abrogated the tumor-promoting effects of HHLA2, highlighting the critical role of c-Met activity in HHLA2-driven HCC (Supplemental Figure 10A).

To expedite tumor formation, we employed a modified approach by co-injecting myr-AKT1, N-RasV12, and c-MET, as establishing HDTVi models solely with N-RasV12 and Met can be time-consuming. This strategy significantly accelerated tumorigenesis. Mice harboring myr-AKT1, N-RasV12, WT c-Met, and HHLA2 displayed a marked increase in tumor burden compared to the control group, as evidenced by the liver-to-body weight ratio. Crucially, substituting WT c-Met with KD c-Met abolished the pro-tumorigenic effects of HHLA2 (Figure 5G). Furthermore, immunohistochemical analysis of livers from the WT c-Met group revealed elevated c-Met phosphorylation and increased CD34+ microvessel density in HHLA2-overexpressing tumors. These changes were significantly attenuated upon replacing WT c-Met with KD c-Met (Figure 5H). These in vivo findings prove that c-Met activation is essential for HHLA2-mediated HCC development and progression.

### c-MET Inhibition Abrogates HHLA2-Driven Hepatocellular Carcinoma (HCC) Progression and Metastasis

Our initial studies investigated the efficacy of c-Met inhibition in suppressing HHLA2-mediated HCC progression using orthotopic xenograft models in nude mice. HepG2-Vec or HepG2-HHLA2 cells were injected into the left liver lobe, followed by alternate-day intraperitoneal administration of the c-Met inhibitor, PHA665752, starting from day three (Figure 6A). Tumor growth and associated pathological changes were monitored over time.

**Figure 6.**
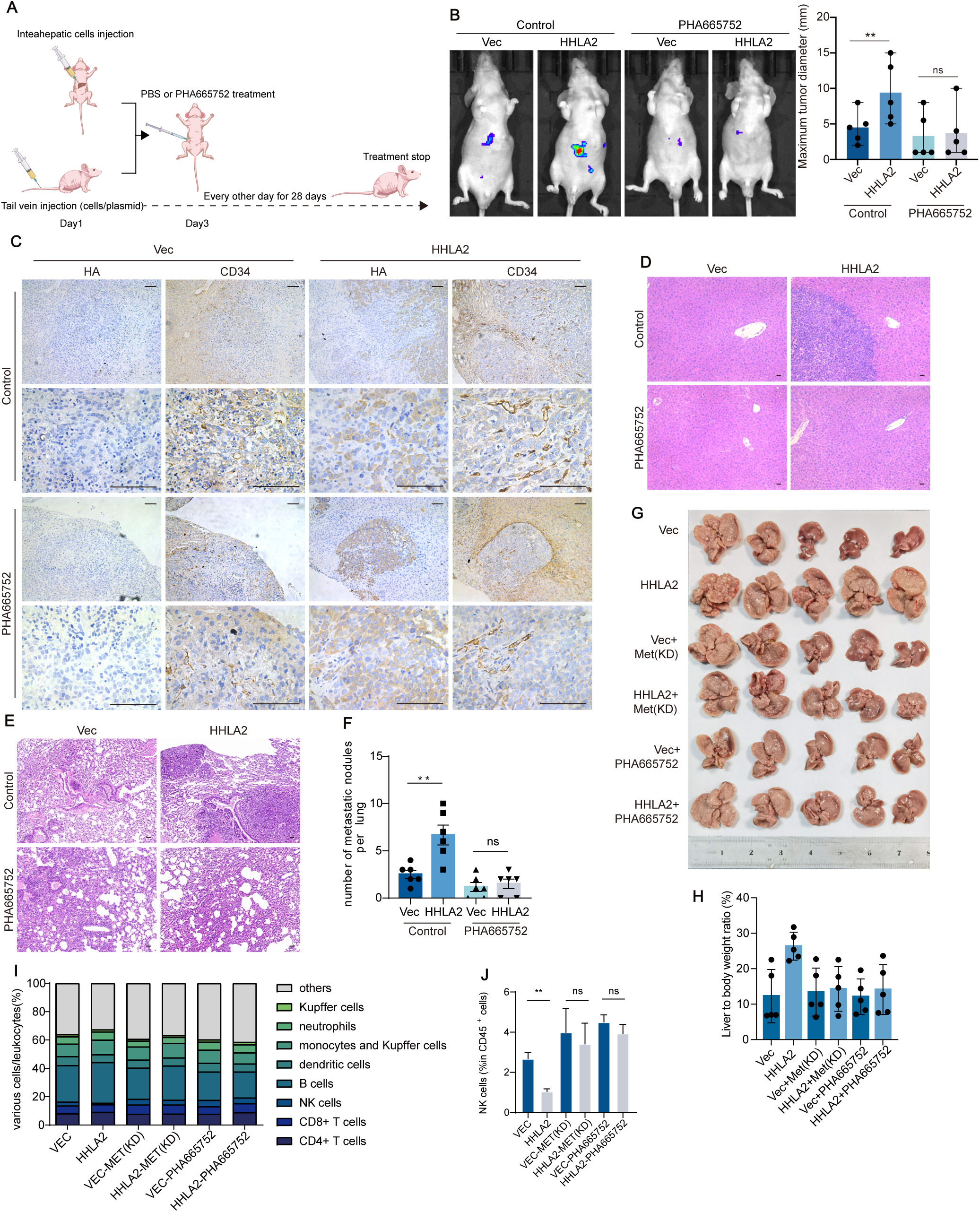
c-Met inhibition abrogates HHLA2-driven HCC progression. **(A)** Schematic representation of intraperitoneal PHA665752 administration. Dosing began on day 3 after HDTVi and continued every other day for 28 days at 20 mg/kg per dose. **(B)** Representative images and quantification of orthotopic HCC xenografts in nude mice. Mice were injected with HepG2-Vec or HepG2-HHLA2 cells, followed by intraperitoneal injection of PHA665752 as outlined in A. The left panel shows representative images of the mice, and the right panel quantifies the largest tumor size (n = 5 mice per group). **(C)** Immunohistochemical staining of xenograft livers for HA (HHLA2) and CD34 (blood vessels). Tissues were obtained from the experiment described in (B). **(D)** H&E staining of uninjected liver lobes to assess HCC metastasis. HepG2-Vec or HepG2-HHLA2 cells were injected into the left liver lobe, followed by PHA665752 treatment as described in (A). **(E, F)** Assessment of HCC lung metastasis in nude mice following tail vein injection of HepG2-Vec or HepG2-HHLA2 cells and PHA665752 treatment. (E) H&E staining of lung tissues. (F) Quantification of tumor nodules. PHA665752 treatment was administered as described in (A) (n = 6 mice per group). **(G–J)** Orthotopic liver tumor development and immune cell infiltration in C57BL/6 mice. (G) Representative images of orthotopic liver tumors following HDTVi of myc-HHLA2 or a control vector, along with N-RasV12/myr-AKT1 and Sleeping Beauty transposon system. (H) Quantification of tumor burden (liver/body weight ratio). (I) Overall proportion of immune cells in different liver tissues. (J) Quantification of NK cell abundance. In this model, either kinase-dead MET (MET-KD) was delivered via HDTVi or PHA665752 was administered as described in A. Data are presented as mean ± SD (n = 5 mice per group). P values were determined by two-tailed Student’s t test. * P <0.05, ** P < 0.01, *** P < 0.001, **** P < 0.0001. Scale bars, 100 μm.

HHLA2 overexpression accelerated intrahepatic tumor growth, as evidenced by increased maximum tumor diameters (Figure 6B, Supplemental Figure 10B), and enhanced tumor angiogenesis, as demonstrated by elevated CD34+ microvascular density (Figure 6C). Importantly, treatment with the c-Met inhibitor PHA665752 effectively countered both the HHLA2-mediated tumor growth and the increase in microvascular density (Figure 6A, 6B, Supplemental Table 3). Furthermore, H&E staining revealed prominent intrahepatic tumor metastasis in un-injected liver lobes from mice injected with HepG2-HHLA2 cells (Figure 6D, Supplemental Table 3). This metastatic spread, a hallmark of advanced HCC with poor prognosis, was entirely abolished by c-Met inhibitor treatment, underscoring its potent anti-metastatic effect.

We established a separate lung metastasis model to strengthen these findings by injecting HepG2 cells via the tail vein in nude mice (Figure 6E, 6F, Supplemental Figure 10C). Consistent with the orthotopic model, mice injected with HHLA2-overexpressing cells displayed significantly higher lung metastatic nodules than the control group. Significantly, treatment with PHA665752 effectively suppressed this enhanced metastatic burden (Figure 6E, 6F), further supporting the anti-metastatic potential of c-Met inhibition.

To further validate these findings and gain mechanistic insights, we utilized a well-established myr-AKT1+ N-RasV12-driven HDTVi HCC model in C57BL/6 mice (Figure 6G, 6H). Consistent with the xenograft model, HHLA2 overexpression significantly increased tumor burden (Figure 6G). Critically, targeting mouse endogenous c-Met with an inhibitor or expressing the dominant-negative KD c-Met effectively abolished this HHLA2-mediated tumor promotion (Figure 6G). The lack of additional benefit observed with PHA665752 treatment in mice with KD c-Met further underscores the pivotal role of c-MET activation by HHLA2 in driving HCC progression.

Previous studies have suggested immunosuppressive functions for HHLA2(31, 32) within the context of HCC. Overexpression of the dominant-negative c-Met mutant effectively abrogated the tumor-promoting effects of HHLA2 (Figure 6G), suggesting that HHLA2’s oncogenic role in HCC is primarily driven by direct enhancement of tumor cell growth rather than broad immunosuppression. To further delineate the potential role of immune cell subsets in HHLA2-mediated HCC progression, we employed flow cytometry to analyze various immune cell populations within liver tissues from HDTVi mice (Supplemental Figure 10D). Notably, neither HHLA2 overexpression nor c-MET inhibition via the dominant-negative mutant or the pharmacological agent PHA665752 significantly impacted the infiltration of B cells, monocytes, dendritic cells, neutrophils, Kupffer cells, or CD4+ and CD8+ T cells (Figure 6I, Supplemental Figure 10E). Furthermore, the activity of both CD4+ and CD8+ T cells remained largely unaffected (Supplemental Figure 10F). Intriguingly, HHLA2 overexpression specifically suppressed the infiltration of natural killer (NK) cells (Figure 6J), a crucial component of the innate immune system with potent anti-tumor activity(31). However, either KD c-Met expression or c-Met inhibition with PHA665752 completely abolished this inhibitory effect. Remarkably, NK cell infiltration in the livers of these mice, regardless of HHLA2 expression, was significantly enhanced and reached comparable levels. While the number of infiltrating NK cells varied between groups, their functional activity remained unchanged (Supplemental Figure 10F). These findings collectively suggest that HHLA2-mediated suppression of NK cell infiltration is mediated through the c-Met pathway in hepatocytes rather than a direct effect on NK cell signaling receptors.

### HHLA2 Promotes c-Met Phosphorylation, Predicts Poor Prognosis, and Shows Promise as a Serum Biomarker

To investigate the clinical relevance of HHLA2 in HCC, we initially assessed its expression and association with c-Met phosphorylation in a cohort of 71 HCC tumor samples obtained from the Shanghai region of China. A robust positive correlation was identified between HHLA2 expression and c-Met phosphorylation (Figure 7A). Specifically, 90.70% of low-HHLA2 tumors exhibited weak c-Met phosphorylation, whereas 64.29% of high-HHLA2 tumors displayed robust c-Met phosphorylation, underscoring a significant association between HHLA2 expression and c-Met activation in HCC (Supplemental Table 4).

**Figure 7.**
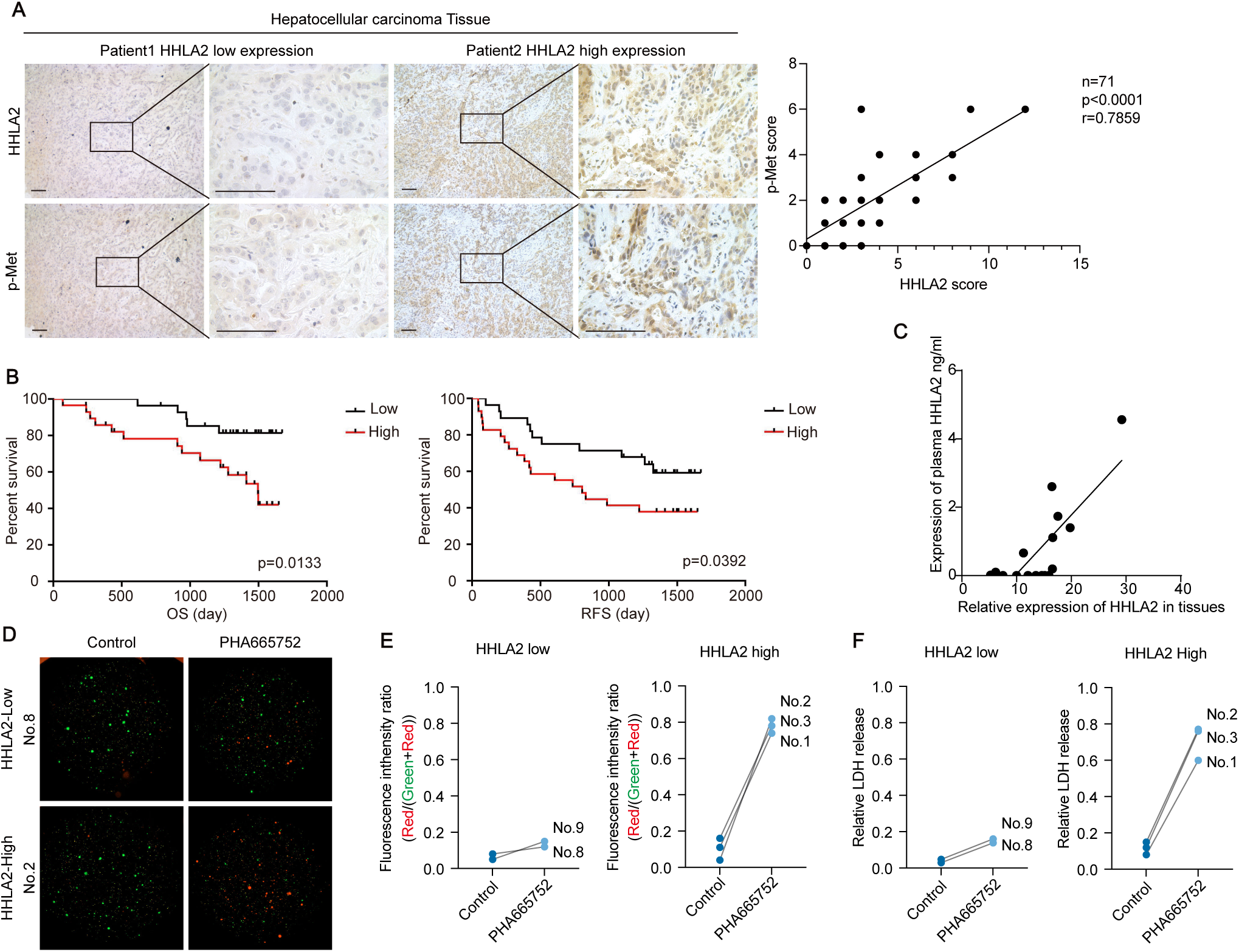
HHLA2 expression correlates with c-Met phosphorylation and predicts efficacy of c-Met inhibitor therapy in HCC. **(A)** Representative images and quantification of p-MET (pY1235) and HHLA2 expression in 71 HCC tissues. Immunohistochemical staining assessed p-MET and HHLA2 expression (left panel). The correlation between p-MET and HHLA2 expression levels was quantified (right panel). Scoring methods are detailed in the Methods section. P value was determined by Pearson correlation analysis. P < 0.0001. **(B)** Kaplan-Meier analysis of OS and recurrence-free survival (RFS) in patients with HCC with high and low HHLA2 expression from the cohort described in (A). P < 0.05. **(C)** Correlation between HHLA2 mRNA expression in HCC tissues and corresponding serum HHLA2 protein levels. x axis: HHLA2 mRNA expression in HCC tissues quantified by qRT-PCR; y axis: serum HHLA2 protein levels quantified by ELISA. **(D–F)** Patient-derived organoid (PDO) response to c-Met inhibition. PDOs derived from HCC tissues in (A), with two exhibiting low HHLA2 expression and three exhibiting high HHLA2 expression, were treated with 15 μM PHA665752 or DMSO for 48 hours. (D) Representative images showing PDO death. (E) Quantification of PDO death. Dead cells are stained red with propidium iodide (PI), and live cells are stained green with Calcein-AM. (F) LDH levels in PDO culture supernatants as a measure of cytotoxicity. Scale bars, 100 μm.

Kaplan-Meier survival analysis, stratifying patients into high and low HHLA2 expression groups based on immunohistochemistry, revealed that the high-HHLA2 group experienced significantly worse overall and progression-free survival (Figure 7B). These findings implicate HHLA2 in promoting HCC progression and negatively influencing patient prognosis, likely through the activation of c-Met signaling.

Furthermore, given the reported utility of other B7 family members, such as PD-L1(32) and B7-H3(33), as circulating biomarkers for cancer detection, we investigated the potential of HHLA2 as a serum biomarker for HCC using liquid biopsies. While current detection methods were insufficiently sensitive to detect HHLA2 in HCC cell line supernatants, we successfully detected HHLA2 in the serum of 13 out of 71 HCC patients exhibiting high HHLA2 tissue expression. Notably, a positive linear correlation was observed between serum and tissue HHLA2 levels (Figure 7C), suggesting the potential utility of HHLA2 as a non-invasive biomarker for HCC. However, further validation and development of more sensitive HHLA2 detection methods are warranted to realize its clinical potential fully.

In conclusion, our study provides compelling evidence that HHLA2 plays a critical role in HCC progression by promoting c-Met phosphorylation. Additionally, HHLA2 holds promise as a potential serum biomarker for HCC, although further investigation is required to confirm its clinical utility.

### HHLA2 Expression Predicts Efficacy of c-Met Inhibitor Therapy in Hepatocellular Carcinoma

To evaluate the potential roles of HHLA2 and c-Met expression in predicting c-Met inhibitor treatment efficacy, we leveraged data from the Cancer Cell Line Encyclopedia (CCLE) database. We extracted transcriptomic, proteomic (RPPA) data and drug sensitivity information for relevant cell lines. C-Met expression was assessed using RPPA data to reflect the relative expression levels of c-Met in tumor cell lines. However, the initial analysis revealed no significant correlation between c-Met expression and c-Met inhibitor efficacy (p>0.05) (Supplemental Figure 11A). Similarly, no correlation was observed between HHLA2 expression, assessed using transcriptomic data, and c-Met inhibitor efficacy (p>0.05) (Supplemental Figure 11B). Further analysis restricted to cell lines with aberrant c-Met expression unveiled a robust positive correlation between HHLA2 expression and c-Met inhibitor efficacy (p=0.0395) (Supplemental Figure 11C). This finding suggested a potential association between HHLA2 expression and c-Met inhibitor sensitivity.

To validate this association in a more clinically relevant setting, we employed patient-derived tumor organoids (PDOs) from five distinct HCC patients (all exhibiting similar c-Met expression levels) (Supplemental Figure 11D, 11E). Notably, PDOs derived from patients with high HHLA2 expression exhibited a more significant response to PHA665752 treatment compared to those with low HHLA2 expression, as evidenced by increased cell death (Figure 7D, 7E, Supplemental Figure 11F) and cytotoxicity (Figure 7F) following treatment. These results further suggest elevated HHLA2 expression may predict a favorable response to c-Met inhibitor therapy in HCC patients.

In conclusion, our data demonstrate that HHLA2 can constitutively activate c-Met, initiating the c-Met signaling pathway and thereby driving the progression of HCC. Furthermore, HHLA2 expression may serve as a prognostic biomarker for a subset of HCC patients with aberrant c-Met expression, enabling the identification and selection of those most likely to benefit from c-Met inhibitor therapy.

## Discussion

Here, we report an oncogenic function of HHLA2 in HCC progression, establishing its direct interaction with and subsequent constitutive activation of c-Met as a crucial driver of HCC development and metastasis. These findings challenge the prevailing understanding of HHLA2 as solely an immune checkpoint molecule (34), highlighting its ability to modulate oncogenic signals independent of its known immunosuppressive functions directly. Our work positions HHLA2 as a potential therapeutic target and prognostic biomarker in HCC, warranting further exploration for its clinical translation.

The oncogenic potential of HHLA2 has been suggested in several cancer types, such as lung cancer and gastric cancer (35–37), where its elevated expression correlates with poor patient prognosis. However, the underlying mechanisms have remained elusive. We provide compelling evidence that HHLA2 directly interacts with c-Met, a receptor tyrosine kinase with established oncogenic function in HCC, through their extracellular domains. This interaction leads to constitutive activation of c-Met, even without its cognate ligand HGF.

HHLA2-mediated c-Met activation deviates from the canonical HGF-induced pathway. HGF binding triggers c-Met internalization and degradation, leading to transient signaling (4). In contrast, HHLA2 stabilizes c-Met at the plasma membrane, preventing its degradation and sustaining downstream signaling, as evidenced by the persistent phosphorylation of c-Met and its effector ERK. This prolonged activation of the c-Met pathway contributes to the sustained proliferation, invasion, and angiogenesis observed in HHLA2-expressing HCC cells. Moreover, HHLA2’s ability to activate c-Met bypasses the requirement for ICAM-1, a known co-receptor for HGF-mediated c-Met activation in HepG2 cells (29). Our findings indicate that HHLA2 can functionally substitute for ICAM-1, resulting in ligand-independent c-Met activation even when ICAM-1 is missing. This suggests a potential mechanism by which HHLA2 might contribute to c-Met activation in HCC tumors with low or absent ICAM-1 expression.

We further demonstrate that N-glycosylation of HHLA2 is essential for its interaction with c-Met and subsequent activation. This finding is consistent with previous studies showing the importance of glycosylation in regulating HHLA2 function in colorectal cancer, as well as the function of other B7 family members (38, 20, 30). This suggests a potential pathway for targeting HHLA2-c-Met interaction through glycosylation modification.

Mechanistically, we identify MMP9 and VEGFA as critical downstream effectors of HHLA2-mediated c-Met activation. HHLA2 upregulates MMP9 and VEGFA expression, promoting HCC invasion and angiogenesis. Notably, inhibition of either MMP9 or VEGFA signaling effectively abrogates the pro-tumorigenic effects of HHLA2, supporting their crucial roles in mediating HHLA2-driven HCC progression.

Our in vitro and in vivo data demonstrate that c-Met activation is indispensable for HHLA2-mediated HCC progression. Pharmacological or genetic inhibition of c-Met effectively abolishes the tumor-promoting effects of HHLA2, including enhanced proliferation, invasion, angiogenesis, and metastasis. This underscores the central role of c-Met signaling in translating the oncogenic potential of HHLA2 in HCC.

While previous studies have implicated HHLA2 in immunosuppression, our findings in the HDTVi HCC mouse model suggest its primary oncogenic mechanism involves direct enhancement of tumor cell growth, rather than broad immune suppression. The absence of HHLA2’s receptors, TMIGD2 and KIR3DL3 (39, 40), on murine immune cells may explain the lack of broad immunosuppression. However, the selective suppression of NK cell infiltration by HHLA2 overexpression, an effect reversed by c-Met inhibition, strongly supports an indirect, c-Met-dependent mechanism for HHLA2-mediated immunosuppression in HCC. Further investigation is required to fully delineate the complex interplay between HHLA2, c-Met signaling, and the immune microenvironment in HCC.

Clinically, we found a strong positive correlation between HHLA2 expression and c-Met phosphorylation in HCC patient samples, suggesting a potential clinical relevance for the HHLA2-c-Met axis. High levels of HHLA2 were also linked to worse patient outcomes, making it a potential biomarker for prognosis. Moreover, we also detected HHLA2 in the blood of HCC patients with high tissue expression, suggesting that it could be developed as a non-invasive serum marker to help detect and monitor the disease.

Finally, our data indicate that HHLA2 expression could be used to predict how patients respond to c-Met inhibitors. In HCC cell lines with high c-Met expression, elevated HHLA2 levels were linked to greater sensitivity to c-Met inhibitors. This was further supported by data from patient-derived tumor organoids, where high HHLA2 expression led to better responses to c-Met inhibitor treatments. More research is needed to confirm this in more extensive clinical trials, but HHLA2 could be a valuable biomarker for personalizing c-Met inhibitor therapy in HCC patients.

In summary, our study establishes HHLA2 as a novel driver of HCC by demonstrating its direct activation of c-Met and consequent tumor promotion. Targeting HHLA2, or its interaction with c-Met, presents a promising therapeutic strategy, potentially overcoming limitations of solely targeting c-Met. Future research clarifying the mechanism of c-Met activation by HHLA2, particularly the role of glycosylation, and validating HHLA2’s biomarker potential will be critical for advancing this therapeutic avenue.

## Methods

### Additional details are available in Supplemental Methods

#### Sex as a biological variable

This study exclusively used male mice due to the significantly higher incidence and more rapid progression of hepatocellular carcinoma (HCC) in male mice compared to females, aligning with human epidemiological trends. While this approach facilitates a more efficient study of HCC development, it limits the generalizability of our findings to female mice.

#### Omics Data Analysis

Publicly available datasets were used to explore HHLA2 expression and its clinical relevance and to investigate the potential link between HHLA2 expression and cancer cell response to therapy. Details of the databases and analysis protocols are provided in the Supplemental Methods.

#### Clinical Samples

Two independent cohorts of HCC tissues were analyzed. Cohort 1 consisted of 176 paired HCC tumor and adjacent non-tumor tissue samples obtained from the Affiliated Cancer Hospital and Institute of Guangzhou Medical University. This cohort was used to analyze HHLA2 expression and its correlation with clinicopathological features. Cohort 2 consisted of 71 HCC tissue samples from the Eastern Hepatobiliary Surgery Hospital in Shanghai, China, and was used to evaluate the correlation between HHLA2 expression and phosphorylated Met (p-Met) levels.

#### Cell Culture

Human HCC cell lines SK-Hep-1, Hep3B, Huh7, MHCC97H, HepG2, SMMC7721, and BEL7402 were cultured as follows: SK-Hep-1, SMMC7721, and BEL7402 in RPMI 1640 medium; Hep3B in Eagle’s Minimum Essential Medium (EMEM); and SK-Hep-1, Hep3B, Huh7, MHCC97H, and HepG2 in Dulbecco’s Modified Eagle Medium (DMEM). All media were supplemented with 10% fetal bovine serum (FBS), except for DMEM used for HUVEC culture, which was supplemented with 20% FBS. Human umbilical vein endothelial cells (HUVECs) were cultured in DMEM supplemented with 20% FBS.

#### Animal Models and In Vivo Experiments

Five-week-old male BALB/c nude mice and seven-week-old male C57BL/6 mice were obtained from the Guangdong Medical Laboratory Animal Center and housed under specific pathogen-free (SPF) conditions.

1. *Mouse Models:* Intrahepatic Xenograft Model: HCC cells were inoculated into the left lateral lobe of the liver of nude mice. 2)Lung Metastasis Model: Lung metastasis was assessed by bioluminescence imaging after tail vein injection of HCC cells into nude mice. 3) Hydrodynamic Co-expression Model: Hydrodynamic tail vein injection was used to deliver various combinations of plasmids encoding HHLA2 (or control), human NRAS, AKT, c-Met (human), and the sleeping beauty transposase (pT3/pCMV-SB) into 8-week-old C57BL/6 mice.
2. *C-Met Inhibitor Treatment:* Mice were treated intraperitoneally with the c-Met inhibitor PHA665752 (20 mg/kg) or vehicle (normal saline) five times per week.
3. *Monitoring and Humane Endpoints:* Mice were monitored for changes in abdominal girth and signs of morbidity or discomfort. Animals were sacrificed at predetermined time points or when tumors reached a maximum size of 2 cm, in accordance with animal welfare guidelines. Further details on sample size determination and allocation are needed in the supplemental methods.

#### Organoid Drug Response Assay

Patient-derived organoids were treated with either c-Met inhibitor PHA665752 (15 μM) or DMSO (vehicle control). After 48 hours, cell viability and apoptosis were assessed using the Calcein-AM/PI Live/Dead Cell Double Staining Kit (Servicebio). Organoids were imaged using a high-content imaging analysis system (AMOview-100, Amoolo Biotech), with Calcein detected at 490/515 nm and propidium iodide (PI) at 535/617 nm. Fluorescence ratios were normalized to vehicle-treated controls. Lactate dehydrogenase (LDH) release in the organoid culture supernatant was measured after 48 hours using the CyQUANT™ LDH Cytotoxicity Assay Kit (Thermo Fisher).

#### Statistics

Statistical analyses were performed using GraphPad Prism version 10. Unpaired or paired Student’s t-tests were used for two-group comparisons. One-way or two-way analysis of variance (ANOVA) followed by appropriate post-hoc tests (e.g., Tukey’s multiple comparisons test) were used for multiple group comparisons. Data normality was assessed using the Shapiro-Wilk test or other appropriate tests. Data are presented as mean ± standard deviation (SD) or median with interquartile range (IQR), as appropriate. P-values < 0.05 were considered statistically significant.

#### Study approval

All patients signed a written informed consent form, agreeing to the use of their data solely for research purposes. This study adhered to the principle outlined in the Declaration of Helsinki (2013 revision) and was approved by the Ethical Review Committees of the Eastern Hepatobiliary Surgery Hospital (No. EHBHKY2020-K-028). All animal experiments were performed according to protocols approved by the Animal Care and Use Committee of Nanjing Drum Tower Hospital and adhered to the Guide for the Care and Use of Laboratory Animals.

## Data availability

Data are available upon reasonable request to the corresponding author. Request for reagents and protocols should be directed to the corresponding author, YW.

## Authors’ contributions

Study concept and design: Yongjie Wei, Feng Shen, Jiahong Wang and Haoshen. Acquisition, analysis and interpretation of data: Xubo Huang, Runya Fang, Yuqian Pang, Zhe Zhang, Jieru Huang, Yingchang Li and Tao Yuan; Manuscript drafting: Xubo Huang, Hao shen and Yongjie Wei; Critical revision of the manuscript for important intellectual content: Silvia Vega-Rubín-de-Celis, Josephine Thinwa; Statistical analysis: Xubo Huang, Yuyi Zeng and Ziying Yao.

XH, YF and YP contributed equally to this work and the author order was determined by seniority.

## Acknowledgements

This work was supported by grants from the National Natural Science Foundation of China (32070718) and the Shenzhen Bay Laboratory Open Fund Project (SZBL2021080601003). We thank Dr. Xiaofei Zhang of Guangzhou Institute of Biomedicine and Health, Chinese Academy of Science for his technical assistance with LS–MS/MS, and Dr. Justin L. Tan of the Shenzhen Bay Laboratory for his valuable suggestions.

Address correspondence to: Yongjie Wei, Guangzhou Medical University, 195 W. Dongfeng Rd, Guangzhou, China, 510182. Phone: +86-22-81340454; Or to: Feng Shen, Department of Hepatic Surgery IV, the Eastern Hepatobiliary Surgery Hospital, 225 Changhai Road, Shanghai 200438, China.; Or to: Jiahong Wang, The affiliate cancer hospital, Guangzhou Medical University, 78 Heng-zhi-gang road, Guangzhou, China, 510095.; Or to: Hao Shen, Department of Hepatic Surgery IV, the Eastern Hepatobiliary Surgery Hospital, 225 Changhai Road, Shanghai 200438.

