## supplemental methods, tables and figures for "Ligand-independent c-Met activation by HHLA2 drives hepatocellular carcinoma and predicts c-MET inhibitor efficacy"

### Omics data analysis

**HHLA2 expression and patient prognosis:** Publicly available datasets were used to explore HHLA2 expression and its clinical relevance.

- HHLA2 protein expression data in normal and tumor tissues were obtained from the Human Protein Atlas (HPA)[46].
- To investigate the association between HHLA2 expression and patient prognosis, we analyzed gene expression and corresponding clinical follow-up data from The Cancer Genome Atlas (TCGA) [The Cancer Genome Atlas: https://portal.gdc.cancer.gov/].
- We further evaluated the relationship between HHLA2 expression or protein phosphorylation levels from The Cancer Proteome Atlas (TCPA) Reverse Phase Protein Array (RPPA) data [The CancerProteome Atlas: https://tcpaportal.org/tcpa/index.html) ].

**HHLA2 expression and drug sensitivity:** The potential link between HHLA2 expression and cancer cell response to therapy was explored using data from the Cancer Cell Line Encyclopedia (CCLE) [http://www.broadinstitute.org/ccle].

- Gene expression (CCLE_DepMap_18Q4_RNAseq_log2 [TPM + 1]), protein expression (CCLE_RPPA_20180123.csv), and drug sensitivity (CCLE_NP24.2009_Drug_-data_2015.02.24) data were downloaded from the CCLE portal.
- Specifically, we analyzed the correlation between HHLA2 and c-Met expression levels with the half-maximal inhibitory concentration (IC50) of c-Met inhibitors.

### Clinical samples

This study utilized two independent cohorts of human HCC tissues. The first cohort comprised 176 HCC samples and was employed to analyze HHLA2 expression and assess its correlation with clinicopathological features. Tissue specimens, including matched tumor and adjacent non-tumor tissues, were obtained from the Affiliated Cancer Hospital and Institute of Guangzhou Medical University. The study protocol was approved by the institution's Ethics Committee above, and written informed consent was obtained from all patients per the Declaration of Helsinki. The second cohort, encompassing 71 HCC samples, was used to evaluate the correlation between HHLA2 expression and p-Met levels. Tissue samples were procured from the Eastern Hepatobiliary Surgery Hospital in Shanghai, China. Approval for this study was granted by the Ethics Committees of both the Affiliated Cancer Hospital of Guangzhou Medical University and the Eastern Hepatobiliary Surgery Hospital, with written informed consent obtained from all participants. All experiments and analyses were conducted with each patient's understanding and written consent.

#### Cell culture

Human HCC cell lines SK-Hep-1, Hep3B, Huh7, MHCC97H, and HepG2 were cultured in Dulbecco's Modified Eagle Medium (DMEM) supplemented with 10% fetal bovine serum (FBS). SK-Hep-1, SMMC7721 and BEL7402 cells were maintained in RPMI 1640 medium containing 10% FBS. Hep3B cells were grown in Eagle's Minimum Essential Medium (EMEM) supplemented with 10% FBS. Human umbilical vein endothelial cells (HUVECs) were cultured in a DMEM medium containing 20% FBS.

#### Quantitative real-time polymerase chain reaction (qRT-PCR)

Total RNA was isolated from HCC cell lines using the FastPure Cell/Tissue Total RNA Isolation Kit (Vazyme, China) and reverse-transcribed into cDNA using the HiScript III RT SuperMix kit (Vazyme, China). qRT-PCR was performed on a CFX96 Touch Real-Time PCR Detection System (Bio-Rad, Hercules, CA, USA) using ChamQ Universal SYBR qPCR Master Mix (Vazyme, China). Specific primers were designed for target genes:

```
HHLA2: 5'-TACAAAGGCAGTGACCATTTGG-3',
       5'-AGGTGTAAATTCCTTCGTCCAGA-3'
MET:   5'-AGCAATGGGGAGTGTAAAGAGG-3';
       5'-CCCAGTCTTGTACTCAGCAAC-3,
MMP9:  5'-TGTACCGCTATGGTTACACTCG-3';
       5'-GGCAGGGACAGTTGCTTCT-3',
MMP2:  5'-TACAGGATCATTGGCTACACACC-3';
       5'-GGTCACATCGCTCCAGACT-3'
MMP7:  5'-GAGTGAGCTACAGTGGGAACA-3';
       5'-CTATGACGCGGGAGTTTAACAT-3'
CDH1:  5'-CGAGAGCTACACGTTCACGG-3';
       5'-GGGTGTCGAGGGAAAAATAGG-3',
CDH2:  5'-TCAGGCGTCTGTAGAGGCTT-3';
       5'-ATGCACATCCTTCGATAAGACTG-3',
VEGFA: 5'-AGGGCAGAATCATCACGAAGT-3';
       5'-AGGGTCTCGATTGGATGGCA-3',
GAPDH: 5'-GGAGCGAGATCCCTCCAAAAT-3';
       5'-GGCTGTTGTCATACTTCTCATGG-3'
```

GAPDH was used as an endogenous reference gene for normalization, and the data were analyzed using the $2^{-\Delta\Delta CT}$ method.

### Western blotting analysis

Cells were lysed in lysis buffers containing a complete protease inhibitor cocktail. Whole-cell lysates were clarified by centrifugation at 12,000 × g for 10 min. Protein concentrations in the supernatants were quantified using a Micro BCA™ Protein Assay Kit (Thermo Scientific, USA). Proteins were then separated by sodium dodecyl sulfate-polyacrylamide gel electrophoresis (SDS-PAGE) using 10% polyacrylamide gels and transferred onto polyvinylidene fluoride (PVDF) membranes (MilliporeSigma, Burlington, MA, USA). Membranes were blocked with 5% non-fat dry milk in Tris-buffered saline with Tween-20 (TBST) for 1-2 h at room temperature. Subsequently, membranes were incubated with primary antibodies diluted in TBST with 5% bovine serum albumin (BSA) overnight at 4 °C. After washing with TBST, membranes were incubated with the appropriate horseradish peroxidase (HRP)-conjugated secondary antibodies in TBST with 5% BSA for 30 min at room temperature. Protein bands were visualized using enhanced chemiluminescence (ECL) detection reagents (refer to the supplementary table for details on specific antibodies used).

### Immunofluorescence (IF) staining

Cell lines: HCC cell lines were fixed with 4% paraformaldehyde, permeabilized with methanol, and subsequently blocked with 4% FBS in phosphate-buffered saline (PBS) for 30 min at room temperature. Cells were then incubated with primary antibodies diluted in PBS with 5% BSA overnight at 4 °C. After washing with PBS, cells were incubated with fluorescence-labeled secondary antibodies diluted in PBS with 5% BSA for 30 min at room temperature. Nuclei were counterstained with 4′,6-diamidino-2-

phenylindole dihydrochloride (DAPI, Beyotime, China) for 5 min. Images were captured using a laser confocal microscope (Zeiss LSM880).

Tissues: Formaldehyde-fixed, paraffin-embedded tissue sections were deparaffinized with xylene, rehydrated through a graded ethanol series, and treated with 0.3% hydrogen peroxide to quench endogenous peroxidase activity. Antigen retrieval was performed by steaming sections in EDTA buffer (pH 8.0) for 15 min. Sections were then incubated with primary antibodies diluted in PBS with 5% BSA overnight at 4 °C. After washing with PBS, sections were incubated with fluorescence-labeled secondary antibodies diluted in PBS with 5% BSA for 30 min at room temperature. Nuclei were counterstained with DAPI (Beyotime, China) for 5 min. IF signals were visualized using a laser confocal microscope (Zeiss LSM880).

### Immunohistochemical (IHC) staining

Formaldehyde-fixed, paraffin-embedded tissue sections were deparaffinized with xylene, rehydrated through a graded ethanol series, and treated with 0.3% hydrogen peroxide to inactivate endogenous peroxidase activity. Antigen retrieval was performed by steaming sections in EDTA buffer (pH 8.0) for 15 min. Sections were then incubated with the primary antibody diluted in PBS with 5% BSA overnight at 4 °C. After washing with PBS, sections were incubated with a horseradish peroxidase (HRP)-conjugated secondary antibody (ZSGB-BIO, China) for 30 min at 37 °C. Immunoreactivity was visualized using DAB chromogenic substrate (Servicebio, China) under microscopic observation. Sections were then counterstained with hematoxylin.

HHLA2 expression scoring: IHC staining intensity and the proportion of positive cells were used to semi-quantitatively evaluate HHLA2 expression. Scores ranged from 0 to 3 based on the following criteria: 0 (0–5% positive cells, low intensity), 1 (6–35% positive cells, low-to-moderate intensity), 2 (36–70% positive cells, moderate intensity), and 3 (>70% positive cells, high intensity). Final expression levels were categorized as low (scores 0–1) or high (scores 2–3). Two experienced pathologists independently assessed the staining in a blinded manner, and mean percentage values were used for analysis.

#### Co-immunoprecipitation (co-IP)

Cells were lysed in appropriate buffers containing a complete protease inhibitor cocktail. Whole-cell lysates were clarified by centrifugation at 12,000 × g for 10 min. For each co-immunoprecipitation reaction, 1 μg of specific antibody and 25 μL of protein A/G magnetic beads (Thermo Scientific, USA) were added to the lysates and incubated overnight at 4 °C with gentle rotation. The beads were then washed extensively with lysis buffer to remove unbound proteins. Finally, immune complexes were eluted from the beads by boiling in SDS-PAGE sample buffer for 5 min at 95 °C.

#### Mass spectrometry (MS) analysis

Following co-immunoprecipitation, the eluted proteins were separated by SDS-PAGE and visualized using silver staining (Beyotime, P0017). For MS analysis, protein bands of interest were excised from the gel and subjected to in-gel trypsin digestion. The resulting peptides were analyzed by a timsTOF Pro mass spectrometer (Bruker) coupled to a NanoElute system (Bruker Daltonics) over a 60-minute gradient. The mass spectrometer was operated in positive ion mode. Raw MS data files were processed using Proteome Discoverer software (version 2.4.0.305; [RRID:SCR_014477]) and the built-in Sequest HT search engine (20) against the UniProt FASTA protein database ( [RRID:SCR_002380]). A maximum of two missed cleavages per peptide was allowed during the database search. Peptide identifications were filtered at a false discovery rate (FDR) of 1%. All other search parameters were set to default.

#### Plasmids, siRNA and cell transfection

Negative control siRNA and siRNAs targeting HHLA2, c-Met, MMP2, and MMP9 were commercially synthesized by Tsingke Biotechnology Company (Guangzhou, China). siRNA transfections were performed using Lipofectamine RNAiMAX transfection reagent (Invitrogen, USA) according to the manufacturer's protocol. All plasmids were transfected into cells using Neofect transfection reagent (NeosBioLab, China) according to the manufacturer's instructions.

### Cell viability assay

Cell viability was assessed using the Cell Counting Kit-8 (CCK-8; Beyotime, China) according to the manufacturer's instructions. Briefly, HCC cells were seeded in 96-well plates at a density of $1.0 \times 10^3$ cells per well in quintuplicate. After incubation for the indicated time points, 10 μL of the CCK-8 solution was added to each well, and the cells were incubated for an additional 1-4 hours following the manufacturer's recommendations. The absorbance at 450 nm was then measured using a Varioskan LUX multimode microplate reader (Thermo Scientific, USA). Higher absorbance values corresponded to greater cell viability.

### Colony formation assays

Two-dimensional (2D) colony formation assay: HCC cells were seeded in triplicate at a density of $1.5 \times 10^3$ cells per well in six-well plates. After two weeks of incubation, cells were fixed with 4% paraformaldehyde for 30 min and stained with 0.5% crystal violet for 1 h. Colonies were visualized and counted under a microscope.

Soft agar colony formation assay: Anchorage-independent growth was assessed using a soft agar colony formation assay. The base layer consisted of 0.6% agar in a complete medium, solidified at room temperature. The top layer contained 0.3% agar in a complete medium supplemented with $5.0 \times 10^3$ HepG2 cells per well. Cells were cultured for two weeks, and colonies were observed under a microscope.

### Cell migration assay

As previously described, a silicone elastomer mask assay was used to quantify HCC cell migration[47]. Briefly, sterile silicone elastomer masks were placed in the center of each well in six-well plates to create a cell-free area. HCC cells ($0.8 \times 10^6$ per well) were then seeded into the wells. After 24 hours of incubation, the masks were removed, and the medium was replaced with a serum-free medium to stimulate cell migration. Cell migration into the formerly cell-free area was monitored and photographed at 0, 24, and 48 hours using an inverted microscope. ImageJ software was used to quantify the area covered by migrated cells at each time point.

### Transwell invasion assay

The invasive potential of HCC cells was assessed using a Transwell invasion assay with Matrigel chambers (Corning, USA). Briefly, HCC cells were seeded in triplicate at a density of $5.0 \times 10^4$ to $1.0 \times 10^5$ cells per well in the upper compartment of the Matrigel invasion chambers. The lower compartment contained 800 μL of DMEM supplemented with 10% FBS and 1 μM PHA665752 as a chemoattractant. After incubation for 24 or 48 hours, cells that invaded through the Matrigel to the lower surface of the membrane were fixed with 4% paraformaldehyde, stained with 0.5% crystal violet, and visualized under a microscope.

### Preparation of conditioned medium:

HCC cells were seeded at a density of $1.0 \times 10^6$ cells per well in six-well plates. To investigate the effects of PHA665752 and si-Met on HCC cell-derived factors, cells were cultured overnight in serum-free medium with or without 1 μM PHA665752 treatment or transfected with either control siRNA or si-Met targeting c-Met. After 24 hours, the conditioned medium was collected and centrifuged to remove cellular debris.

The resulting supernatant was then used for subsequent experiments, including ELISA and HUVEC tube formation assays.

### HUVEC tube formation assay

The angiogenic potential of the conditioned medium was assessed using a HUVEC tube formation assay. Briefly, 96-well plates were pre-cooled and coated with 50 µL/well of Matrigel (Corning, USA). The Matrigel was allowed to solidify at 37 °C for 30 min. HUVECs were harvested using trypsin and resuspended in a serum-free medium. Aliquots of 100 µL of cell suspension (containing $2.0 \times 10^4$ cells/well) were then plated onto the solidified Matrigel. The plates were incubated for 6 hours at 37 °C in a humidified incubator containing 5% $CO_2$. Five images were captured per well using an inverted microscope. ImageJ software quantified the number of tube-like structures formed by the HUVECs. Experiments were performed in triplicate and independently repeated three times.

### RNA sequencing analysis

Total RNA was isolated from HepG2 cells treated with control medium (n = 3) or medium overexpressing HHLA2 (OE-HHLA2; n = 3) using established protocols. RNA sequencing libraries were then constructed and sequenced on the Illumina NovaSeq6000 (Biomarker Technologies sequencing platform) using paired-end 150 bp reads (PE150). Briefly, polyadenylated mRNA was enriched from total RNA using oligo(dT) magnetic beads. The isolated mRNA was fragmented, reverse transcribed into cDNA, and purified. The fragmented cDNA was end-repaired, A-tailed, and ligated with sequencing adapters. Following library size selection and quality control, libraries were sequenced using the Illumina NovaSeq6000 platform. Sequencing data were analyzed using a standard pipeline, including quality control, read alignment to the human genome, gene quantification, and differential expression analysis. Genes were considered differentially expressed if they exhibited a |log2(fold change)| exceeding 0.4 and a p-value less than 0.05.

#### VEGFA Enzyme-linked immunosorbent assay (ELISA)

Soluble VEGFA protein levels in cell culture supernatants were quantified using a commercially available Human VEGF ELISA Kit (Beyotime, China) according to the manufacturer's protocol. Briefly, samples and standards were incubated in ELISA plates pre-coated with capture antibodies specific for VEGFA. After washing steps to remove unbound substances, a horseradish peroxidase (HRP)-conjugated detection antibody was added. The amount of bound HRP-conjugated antibody was then measured using a chromogenic substrate with absorbance measured at 450 nm. The concentration of VEGFA in the samples was extrapolated from a standard curve generated using known concentrations of recombinant human VEGF.

#### Split luciferase complementation assay:

**Stable cell lines generation:** LgBiT fragment of NanoLuc Luciferase（Promega, UK, N2681） and a flexible linker (LgBiT-GSSGGGGSGGGGSSG-) was insert behind the signal peptide (sp) sequences of full-length HHLA2 (between amino acid 23-24) or MET (between amino acid 25-26) using Gibson assembly (NEB, UK, M0530S). The resulting sp-LgBiT-Linker-HHLA2 and sp-LgBiT-Linker-MET open reading frames were then cloned into lentiviral expression vector plenti-IRES-Puro (Origene, USA, PS100069). Lentiviral particles were produced according to the manufacturer's instructions and used to transduce HEK293 cells. Following selection with puromycin (1 μg/mL), stable cell lines exhibiting surface expression of LgBiT-HHLA2 or LgBiT-MET were established.

**Recombinant protein expression and purification:** The extracellular domains (ExD) of HHLA2 or c-MET were fused with the SmBiT fragment of NanoLuc Luciferase （Promega, UK, N2681） and a 6XHis tag at the c-terminus. The resulting constructs were then inserted into the BacMam-pCMV-DEST vector from the ViraPower BacMam Expression system (Thermo Scientific，USA, A24233 PMID: 25299155). BacMam virus particles containing the HHLA2ExD-SmBit-6His or c-MET-ExD-SmBit-6His expression frames were produced in Sf9 cells following the manufacturer's instructions and subsequently used to infect suspension-cultured HEK293S cells. After 3 days, the cell culture supernatant was passed through a Ni-NTA column (Thermo Scientific，USA， K950-1) to purify the secreted recombinant proteins HHLA2ExD-SmBit-6His or c-METExD-SmBit-6His.

**Luciferase Assay:** $1x10^4$ 293-LgBiT-HHLA2, or 293-LgBiT-MET cells were seeded in 96-well plates and cultured overnight. Recombinant c-MET-ExD-SmBit-6His or HHLA2ExD-SmBit-6His proteins were added to the respective wells at concentrations ranging from 1 μg to 30 μg, followed by 1 hour incubation. Subsequently, Nano-Glo Live Cell Substrate was added according to the manufacturer's instructions (Promega, UK). Luminescence was measured after a 15-minute incubation period.

### Protein in vitro binding assay

The in vitro interaction between HHLA2 and c-met was examined using two methods. In the first method, 20 μg purified HHLA2ExD-SmBit-6His and c-MET-ExD-SmBit-6His recombinant proteins were incubated with anti-HHLA2-agarose beads at 4°C overnight. After washing three times with pre-cold PBS buffer to remove unbound proteins, the presence of c-MET was detected using an anti-c-MET antibody. In the second method, 1 μg/well of purified c-MET-ExD-SmBit-6His protein was cross-linked overnight to the bottom of a 96-well plate. Then, 1 μg to 20 μg of HHLA2ExD-SmBit-6His protein was added to each well and incubated for one hour. After three washes with PBS to remove unbound HHLA2, the amount of HHLA2 bound to c-met in the wells was quantified using an ELISA kit (CwBio, China, CW0050).

### Animal models and in vivo experiments

All animal experiments were performed according to protocols approved by the Animal Care and Use Committee of Nanjing Drum Tower Hospital and adhered to the Guide for the Care and Use of Laboratory Animals.

#### Mice

Five-week-old male BALB/c nude mice and seven-week-old male C57BL/6 mice were purchased from Guangdong Medical Laboratory Animal Center and housed in specific pathogen-free facilities.

#### In vivo models

Intrahepatic xenograft models**:** Stable HCC cell lines (HepG2-vec and HepG2-HHLA2) were established and inoculated into the left lateral lobe of the liver (2 million cells per mouse) of nude mice. Tumor growth was monitored by bioluminescence imaging (IVIS Lumina XRMS, PerkinElmer) four weeks after inoculation. Mice were then sacrificed, and livers were harvested, weighed, imaged, and processed for further analysis. Specifically, a portion of the tumor tissue and adjacent liver lobe were fixed in 4% paraformaldehyde for hematoxylin and eosin (H&E) and immunohistochemistry (IHC) staining, while the remaining tissues were snap-frozen in liquid nitrogen for subsequent analysis.

Lung metastasis models**:** Nude mice (n = 6 per group) were injected with stable HCC cells (HepG2-vec and HepG2-HHLA2) via the tail vein (1 million cells per mouse). Lung metastasis was assessed by bioluminescence imaging nine weeks after injection.

Mice were then sacrificed, and lungs were harvested, imaged, and fixed in 4% paraformaldehyde for H&E and IHC staining. The remaining lung tissues were snap-frozen in liquid nitrogen for further analysis.

Hydrodynamic co-expression models: To model NRAS/AKT-driven or AKT/c-Met-driven HCC, plasmids encoding pT3-HHLA2 (or control), pT3-NRAS (human), pT3-AKT, pCMV/SB, pT3-c-Met (human), or combinations thereof, were hydrodynamically injected into the tail vein of eight-week-old C57BL/6 mice (dosages specified in the text). For the NRAS/AKT/c-Met model, wild-type or dominant-negative c-Met plasmids were used.

C-Met inhibitor treatment: Mice were treated with c-Met inhibitor PHA665752 (20 mg/kg) or vehicle (normal saline) intraperitoneally five times per week.

Monitoring and humane endpoints: All mice were monitored for abdominal girth and signs of morbidity or discomfort throughout the experiment. Mice were sacrificed at designated time points, with a maximum tumor size not exceeding 2 cm according to animal welfare guidelines. Sample sizes were determined based on preliminary data indicating group differences. No statistical power calculations were performed to predetermine sample size.

### Tissue dissociation and organoid culture

Fresh liver tumor tissue was obtained from patients undergoing hepatectomy for HCC. Blood, fat, necrotic tissue, and connective tissue were meticulously removed to enrich tumor cells. The remaining tissue was then dissected into small pieces. The tissue fragments were washed in 5 mL of 5 mM PBS/EDTA for 15 minutes at room temperature to remove residual blood. Subsequently, the tissue was incubated in 5 mL of 1 mM PBS/EDTA containing 2x TrypLE for one hour at 37 °C to promote enzymatic dissociation. The dissociated cell clusters were liberated from the remaining tissue fragments by applying gentle mechanical force with a pipette. The resulting cell suspension was centrifuged at 350 g to pellet the cells. The cell pellet was then resuspended in a mixture of 50% Matrigel and organoid culture medium at a density of $2X10^5$/mL. Aliquots of 100 μL of the cell suspension were plated onto a 24-well plate. The plates were incubated for 30 minutes at 37 °C and 5% $CO_2$ for Matrigel solidification. Following solidification, 1 mL of fresh organoid culture medium was added to each well. The culture medium was replaced every 72 hours.

### Organoid drug response assay

The c-Met inhibitor PHA665752 (15 μM, HY-15735, MCE) or dimethyl sulfoxide (DMSO, vehicle control) was added to the PDOs. After 48 hours, cell viability and apoptosis were assessed using the Calcein-AM/PI Live/Dead Cell Double Staining Kit (G1707, Servicebio) according to the manufacturer's instructions. Stained organoids were imaged with a high-content imaging analysis system (AMOview-100, Amoolo Biotech), detecting Calcein at 490/515 nm and PI at 535/617 nm. Fluorescence ratios indicating cell viability and apoptosis were normalized to vehicle-treated controls.Organoid culture supernatant was collected 48 hours post-treatment, and cell damage was evaluated using the CyQUANT™ LDH Cytotoxicity Assay Kit (C20300, Thermo Fisher) as per the manufacturer's protocol.

### Flow cytometry analysis of immune cell infiltrates

Liver tissues were mechanically dissociated using a mortar and pestle to isolate infiltrating immune cells. Single-cell suspensions were washed with PBS containing 1% FBS and red blood cells were lysed using an appropriate lysis buffer. Cells were then stained with fluorochrome-conjugated antibodies specific for various immune cell surface markers in the same buffer. The following antibodies were used:

- Anti-human CD45 (to identify human leukocytes)
- Anti-mouse CD19 (B cells)
- Anti-mouse CD11c+ and MHCII+ (dendritic cells)
- Anti-mouse CD11b+ (monocytes and Kupffer cells)
- Anti-mouse CD11b+ and F4/80+ (Kupffer cells)
- Anti-mouse CD11b+ and Ly6G+ (neutrophils)
- Anti-mouse NK1.1 (NK cells)
- Anti-mouse CD107a+ and NK1.1+ (activated NK cells)
- Anti-mouse CD8+ (CD8+ T cells)
- Anti-mouse CD69+ and CD8+ (activated CD8+ T cells)
- Anti-mouse CD4+ (CD4+ T cells)
- Anti-mouse CD69+ and CD4+ (activated CD4+ T cells)

Flow cytometry was analyzed using a BD FACSAria™ III Cell Sorter (BD Bioscience, USA). Data were analyzed using FlowJo software v.10.

### Statistical analysis

Statistical analyses were performed using GraphPad Prism version 10 (La Jolla, CA, USA). For comparisons between two groups, unpaired or paired Student's t-tests were used as appropriate based on experimental design. For comparisons between multiple groups, one-way or two-way analysis of variance (ANOVA) followed by post-hoc tests (e.g., Tukey's multiple comparisons test) were used. Normality of data distribution was assessed using appropriate tests (e.g., the Shapiro-Wilk test). Data were presented as mean ± standard deviation (SD) or median with interquartile range (IQR) depending on the distribution of the data. P-values less than 0.05 ($p < 0.05$) were considered statistically significant.

### Supplemental Tables

**Supplemental Table 1.** Correlation between HHLA2 Expression and Clinicopathological Factors in 176 HCC Patients from Guangdong, China

| Characteristics | No. of patients | HHLA2 expression (%) | | *p*-value |
|---|---|---|---|---|
| | | Low | High | |
| Gender | | | | |
| Female | 21 | 7 (33.3%) | 14 (66.7%) | |
| Male | 155 | 76 (49.0%) | 79 (51.0%) | 0.176 |
| Age (years) | | | | |
| ≤ 50 | 93 | 38 (40.9%) | 55 (59.1%) | |
| > 50 | 83 | 45 (54.2%) | 38 (45.8%) | 0.076 |
| AFP (ng/ml) | | | | |
| ≤ 400 | 101 | 47 (46.5%) | 54 (53.5%) | |
| > 400 | 75 | 36 (48.0%) | 39 (52.0%) | 0.847 |
| HBsAg | | | | |
| Negative | 11 | 6 (54.5%) | 5 (45.5%) | |
| Positive | 165 | 77 (46.7%) | 88 (53.3%) | 0.612 |
| Liver cirrhosis | | | | |
| No | 37 | 21 (56.8%) | 16 (43.2%) | |
| Yes | 139 | 62 (44.6%) | 77 (55.4%) | 0.188 |
| Tumor size (cm) | | | | |
| ≤ 5 | 102 | 62 (60.8%) | 40 (39.2%) | |
| > 5 | 74 | 21 (28.4%) | 53 (71.6%) | **< 0.001** |
| Tumor number * | | | | |
| Single | 164 | 77 (47.0%) | 87 (53.0%) | |
| Multiple | 12 | 6 (50.0%) | 6 (50.0%) | 0.838 |
| Satellite nodule | | | | |
| No | 155 | 75 (48.4%) | 80 (51.6%) | |
| Yes | 21 | 8 (38.1%) | 13 (61.9%) | 0.375 |
| Tumor capsule | | | | |
| No/incomplete | 105 | 45 (42.9%) | 60 (57.1%) | |
| Complete | 71 | 38 (53.5%) | 33 (46.5%) | 0.164 |
| Tumor differentiation | | | | |
| I-II | 125 | 56 (44.8%) | 69 (55.2%) | |
| III-IV | 51 | 27 (52.9%) | 24 (47.1%) | 0.326 |
| Vascular invasion | | | | |
| No | 157 | 79 (50.3%) | 78 (49.7%) | |
| Yes | 19 | 4 (21.1%) | 15 (78.9%) | **0.016** |
| BCLC stage | | | | |
| 0 | 16 | 10 (62.5%) | 6 (37.5%) | |
| A | 80 | 50 (62.5%) | 30 (37.5%) | |
| B | 62 | 17 (27.4%) | 45 (72.6%) | |
| C | 18 | 6 (33.3%) | 12 (66.7%) | **< 0.001** |
| Recurrence or metastasis | | | | |
| Negative | 80 | 54 (67.5%) | 26 (32.5%) | |
| Positive | 96 | 29 (30.2%) | 67 (69.8%) | **< 0.001** |

* Single or multiple HCC: either one tumor or two and above tumors within the liver

**Supplemental Table 2.** Univariate and Multivariate Analyses of Postoperative Prognosis in 176 HCC Patients

| | OS | | | | TTR | | | |
|---|---|---|---|---|---|---|---|---|
| | Univariate | Multivariate | | | Univariate | Multivariate | | |
| Variables* | *p*-value | *p*-value | HR | 95% CI | *p*-value | *P*-value | HR | 95% CI |
| Gender (Female vs. Male) | NS | NS | | | NS | NS | | |
| Age, years (≤ 50 vs. > 50) | NS | NS | | | NS | NS | | |
| AFP (ng/mL) (≤ 400 vs. > 400) | < 0.001 | < 0.001 | 2.539 | 1.623-3.973 | < 0.001 | < 0.001 | 2.756 | 1.794-4.235 |
| HBsAg (Negative vs. Positive) | NS | NS | | | NS | NS | | |
| Liver cirrhosis (No vs. Yes) | 0.003 | 0.009 | 2.586 | 1.272-5.257 | 0.003 | 0.021 | 2.142 | 1.124-4.083 |
| Tumor size (cm) (≤ 5 vs. > 5) | < 0.001 | NS | | | < 0.001 | NS | | |
| Tumor number (Single vs. Multiple) | NS | NS | | | NS | NS | | |
| Satellite nodule (No vs. Yes) | NS | NS | | | 0.029 | NS | | |
| Tumor capsule (No/ incomplete vs. Complete) | NS | NS | | | NS | NS | | |
| Tumor differentiation (I-II vs. III-IV) | NS | NS | | | NS | NS | | |
| Vascular invasion (No vs. Yes) | < 0.001 | < 0.001 | 5.169 | 2.792-9.569 | < 0.001 | < 0.001 | 4.673 | 2.630-8.304 |
| HHLA2 (Low *versus* High) | < 0.001 | 0.004 | 2.093 | 1.261-3.475 | < 0.001 | < 0.001 | 2.758 | 1.719-4.426 |

**Supplemental Table 3** Number and Proportion of Intrahepatic Metastases in 4 Groups of Orthotopic Liver xenograft Model.

| | Intrahepatic metastasis |
|---|---|
| HepG2-vec | 0/5 |
| HepG2-HHLA2 | 2/5 |
| HepG2-vec-PHA665752 | 0/5 |
| HepG2-HHLA2-PHA665752 | 0/5 |

**Supplemental Table 4.** Correlation between HHLA2 Expression and p-MET levels of a cohort of 71 HCC patients from Shanghai, China

| | | NO. expression of p-MET(%) | | | |
|---|---|---|---|---|---|
| | | -/+ | ++/+++ | Total | P-value |
| | -/+ | 39(54.93%) | 4(5.63%) | 43(60.56%) | |
| HHLA2 | ++/+++ | 10(14.08%) | 18(25.36%) | 28(39.44%) | |
| | Total | 49(69.01%) | 22(30.99%) | 71(100%) | P<0.0001 |

Fig. S1

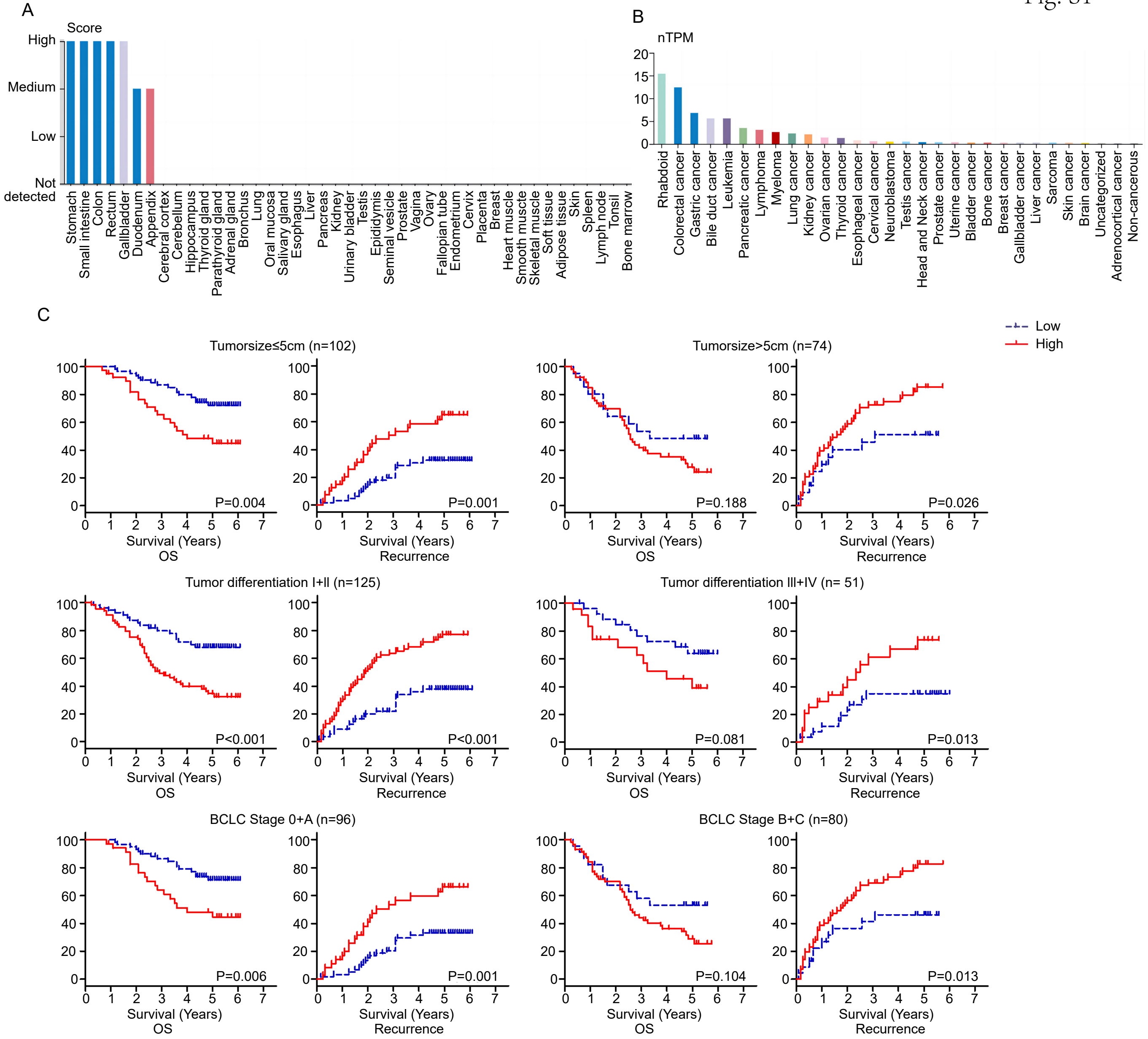


**Supplemental Figure 1. HHLA2 expression correlates with survival in patients with HCC. (A, B)** HHLA2 mRNA expression in normal human tissues (A) and cancer cell lines (B). Data are presented as relative values (A) and transcripts per million (TPM) (B). Data sources: Human Protein Atlas and The Cancer Genome Atlas (TCGA). P values were determined by unpaired two-tailed t test. **(C)** Kaplan-Meier survival curves for overall survival and recurrence in the HCC cohort, stratified by tumor size (≤5 cm or >5 cm), tumor differentiation grade, and Barcelona Clinic Liver Cancer (BCLC) stage. P values were determined by log-rank test. * P < 0.05, ** P < 0.01, *** P < 0.001, **** P < 0.0001.

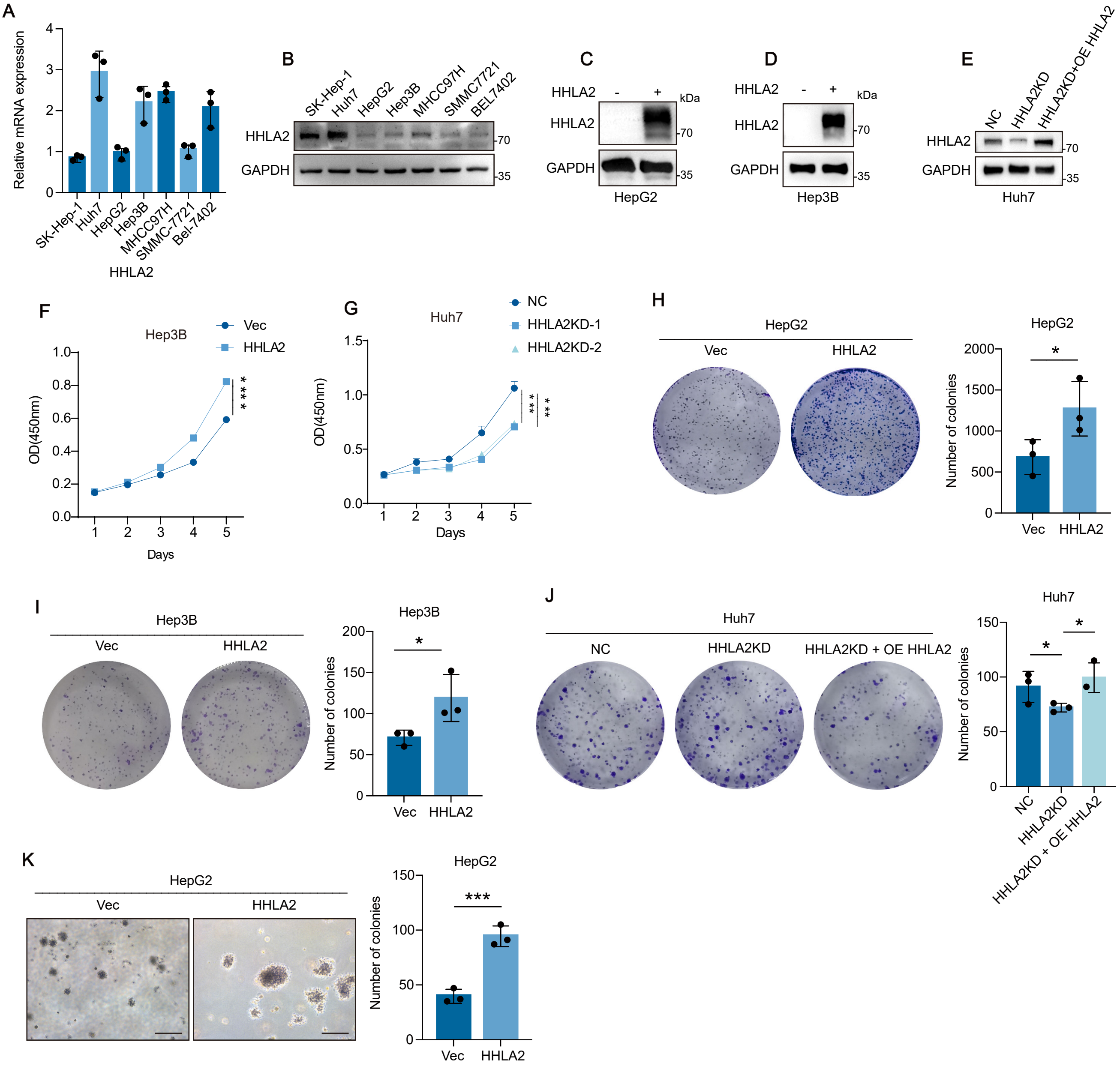


**Supplemental Figure 2. HHLA2 promotes HCC cell growth in vitro. (A, B)** HHLA2 mRNA (A) and protein (B) expression in HCC cell lines. **(C-E)** Western blot validation of HHLA2 overexpression in HepG2 (C) and Hep3B (D) cells and HHLA2 knockdown in Huh7 (E) cells. **(F, G)** Cell proliferation assessed by CCK-8 assay in Hep3B (F) and Huh7 (G) cells with HHLA2 overexpression or knockdown. P values were determined by one-way ANOVA. **(H-J)** Two-dimensional colony formation assays in HepG2 (H), Hep3B (I), and Huh7 (J) cells with HHLA2 overexpression or knockdown (2 weeks). **(K)** Soft agar colony formation assay in HepG2-Vec and HepG2-HHLA2 cells. P values were determined by two-tailed Student's t test. * $P < 0.05$, ** $P < 0.01$, *** $P < 0.001$, **** $P < 0.0001$. Scale bars, 100 μm.

Fig. S3

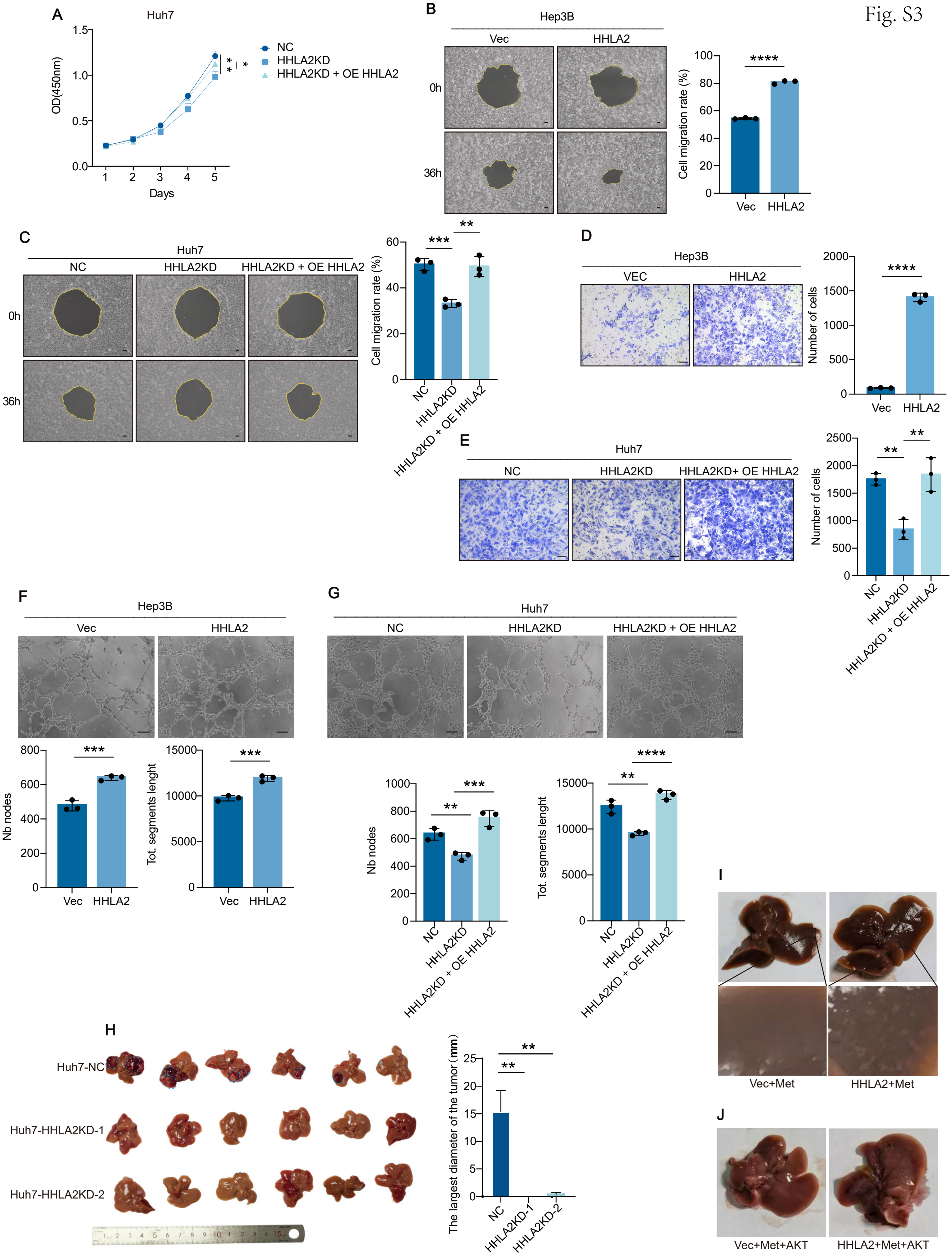
A
Huh7
OD(450nm)
Days
NC
HHLA2KD
HHLA2KD + OE HHLA2
B
Hep3B
Vec
HHLA2
0h
36h
Cell migration rate (%)
C
Huh7
NC
HHLA2KD
HHLA2KD + OE HHLA2
D
Hep3B
VEC
HHLA2
Number of cells
E
Huh7
HHLA2KD+ OE HHLA2
F
Nb nodes
Tot. segments lenght
G
H
Huh7-NC
Huh7-HHLA2KD-1
Huh7-HHLA2KD-2
The largest diameter of the tumor (mm)
HHLA2KD-1
HHLA2KD-2
I
Vec+Met
HHLA2+Met
J
Vec+Met+AKT
HHLA2+Met+AKT

**Supplemental Figure 3. HHLA2 promotes aggressive phenotypes in HCC. (A)** Cell proliferation assessed by CCK-8 assay in the indicated Huh7 cells. **(B, C)** Cell migration assays in Hep3B (B) and Huh7 (C) cells with HHLA2 overexpression or knockdown. **(D, E)** Transwell invasion assays in Hep3B (D) and Huh7 (E) cells with HHLA2 overexpression or knockdown. **(F, G)** HUVEC tube formation assays using conditioned media from Hep3B (F) or Huh7 (G) cells with HHLA2 overexpression or knockdown. **(H)** Representative images and quantification of liver tumor sizes in orthotopic xenografts derived from Huh7 cells expressing either a nontargeting shRNA (Huh7-shNC) or an shRNA targeting HHLA2 (Huh7-HHLA2KD). n = 6 mice per group. **(I, J)** Representative images of orthotopic liver tumors in C57BL/6 mice. Tumors were induced by hydrodynamic tail vein injection (HDTVi) of the following combinations: (I) HHLA2 or a control vector, along with c-Met and Sleeping Beauty transposon system; (J) HHLA2 or a control vector, c-Met, myr-AKT1 (constitutively active AKT1), and Sleeping Beauty transposon system. P values were determined by one-way ANOVA (A) or two-tailed Student’s t test (B–J). * $P < 0.05$, ** $P < 0.01$, *** $P < 0.001$, **** $P < 0.0001$. Scale bars, 100 μm.

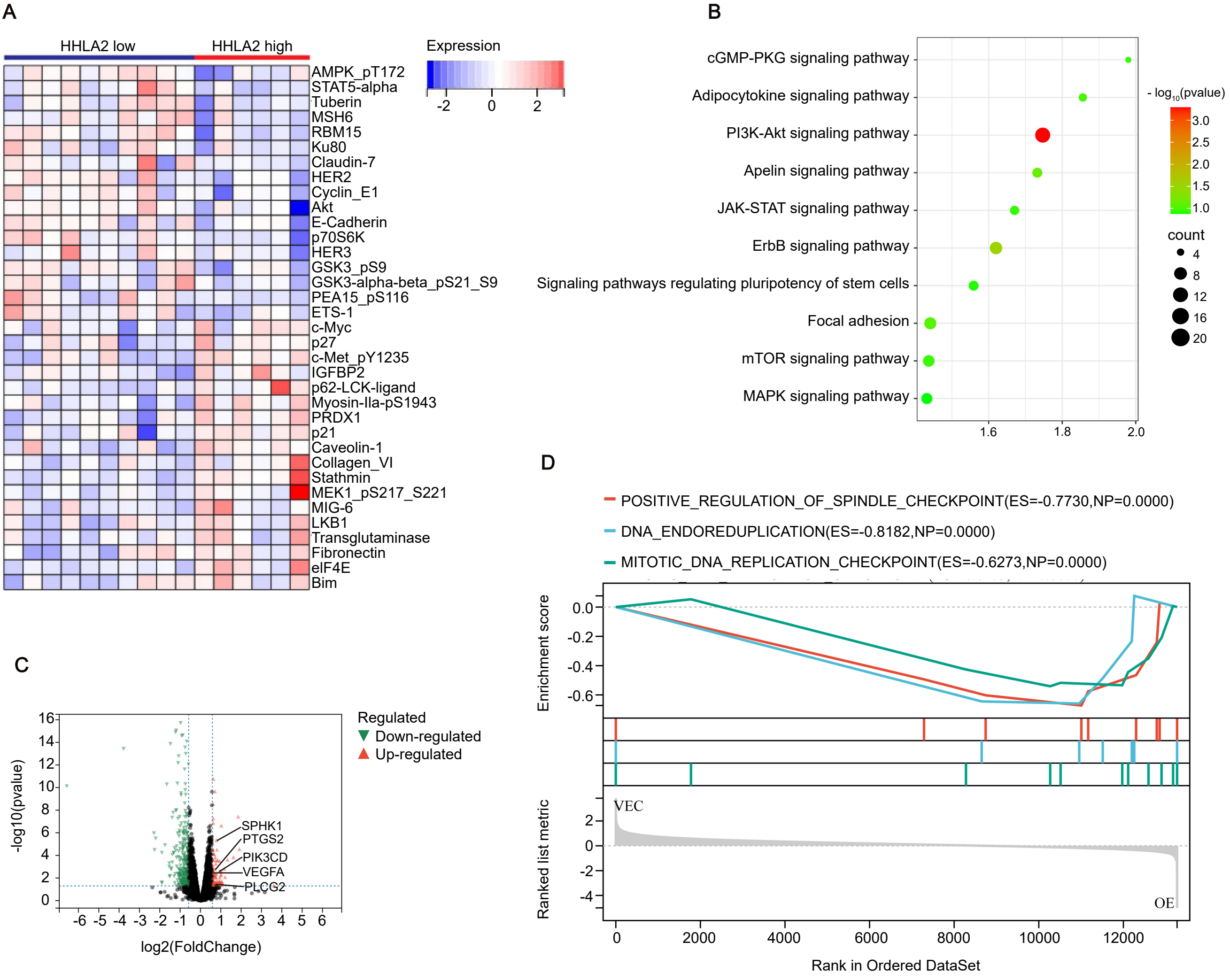


**Supplemental Figure 4. HHLA2 expression modulates specific proteins and pathways. (A)** Heatmap analysis of differentially expressed proteins and phosphoproteins in HCC samples with high or low HHLA2 expression from TCGA and The Cancer Proteome Atlas (TCPA) databases. The heatmap displays proteins and phosphoproteins with a fold change greater than 1.2 between high and low HHLA2–expressing HCC samples. **(B)** Kyoto Encyclopedia of Genes and Genomes (KEGG) pathway enrichment analysis of differentially expressed proteins identified in (A). Only pathways with $P < 0.05$ are shown. **(C)** Volcano plot of differentially expressed mRNAs in HepG2 cells following HHLA2 overexpression. Genes with |log2 (fold change) | > 0.4 and $P < 0.05$ are highlighted (n = 3 samples per group). **(D)** Gene set enrichment analysis (GSEA) of cell proliferation processes. GSEA was performed on the differentially expressed genes identified in (C). NES, normalized enrichment score.

Fig. S5

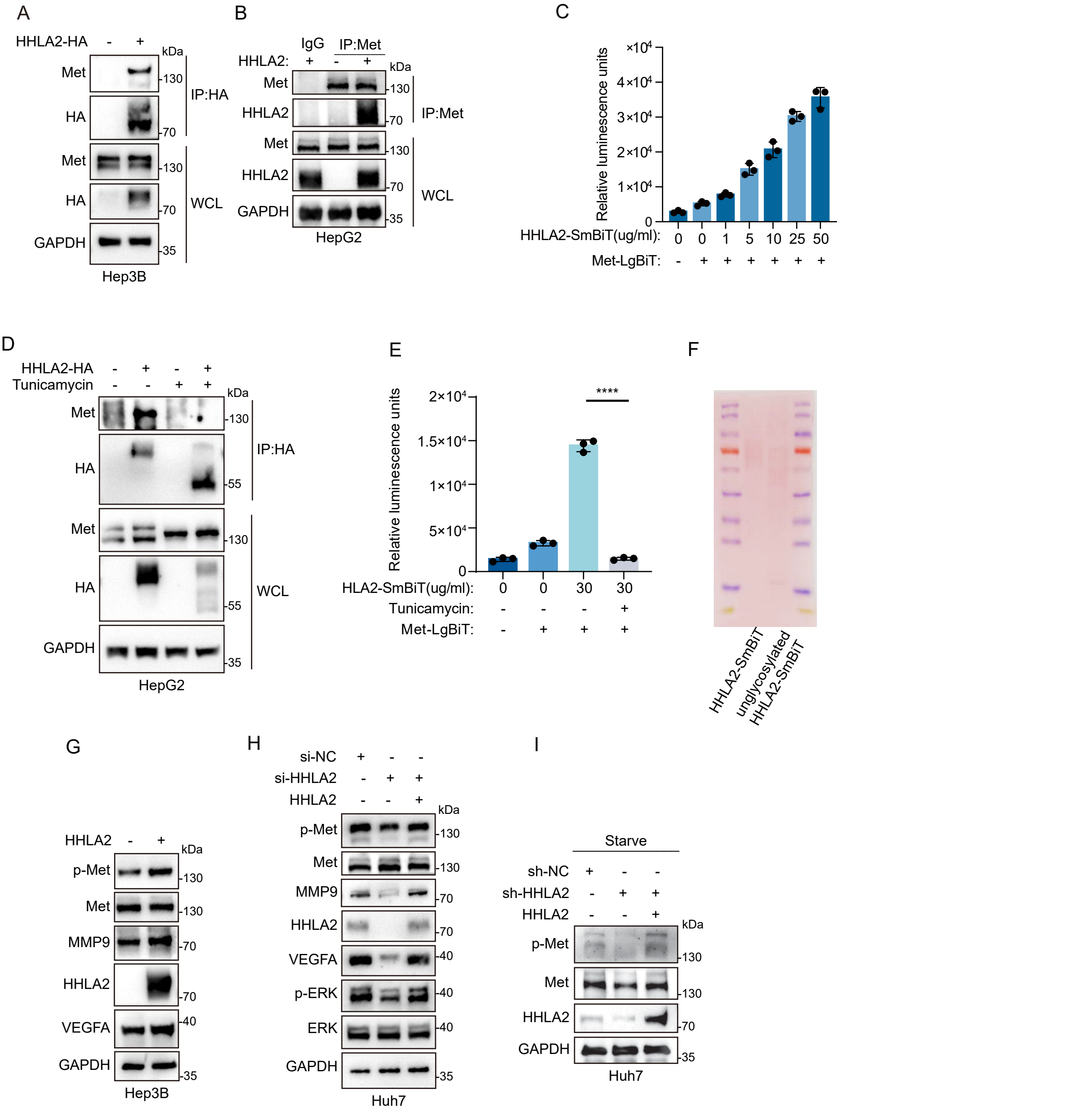


**Supplemental Figure 5. HHLA2 interacts directly with c-Met. (A)** Coimmunoprecipitation of overexpressed HHLA2-HA and endogenous c-MET in Hep3B cells. **(B)** Coimmunoprecipitation of endogenous c-MET and HHLA2 in HepG2 cells. The antibodies used for immunoprecipitation (IP) and Western blot are indicated. **(C)** Split luciferase complementation assay to define the interaction domains between HHLA2 and c-Met. Luminescence was measured following the addition of indicated amounts of recombinant HHLA2-ExD-SmBit-6His to 293-LgBiT-MET cells. **(D)** Coimmunoprecipitation of HHLA2-HA with endogenous c-Met in HepG2-Vec or HepG2-HHLA2 cells treated with 1 µg/ml tunicamycin (N-linked glycosylation inhibitor) or DMSO (control) for 24 hours. **(E, F)** In vitro interaction between N-linked glycosylated or unglycosylated HHLA2 and c-MET assessed by split luciferase complementation assay. Luminescence was measured following the addition of indicated concentrations of glycosylated or unglycosylated HHLA2-ExD-SmBit-6His to 293-LgBiT-HHLA2 or 293-LgBiT-MET cells, respectively (E). P values were determined by two-tailed Student's t test. **** P < 0.0001. Ponceau S staining of input HHLA2-ExD-SmBit-6His proteins is shown in F. **(G-I)** Western blot analysis of c-MET signaling pathway proteins in Hep3B cells with or without HHLA2 overexpression (G) and Huh7 cells with or without HHLA2 knockdown and rescue under normal (H) or starvation (I) conditions.

Fig. S6

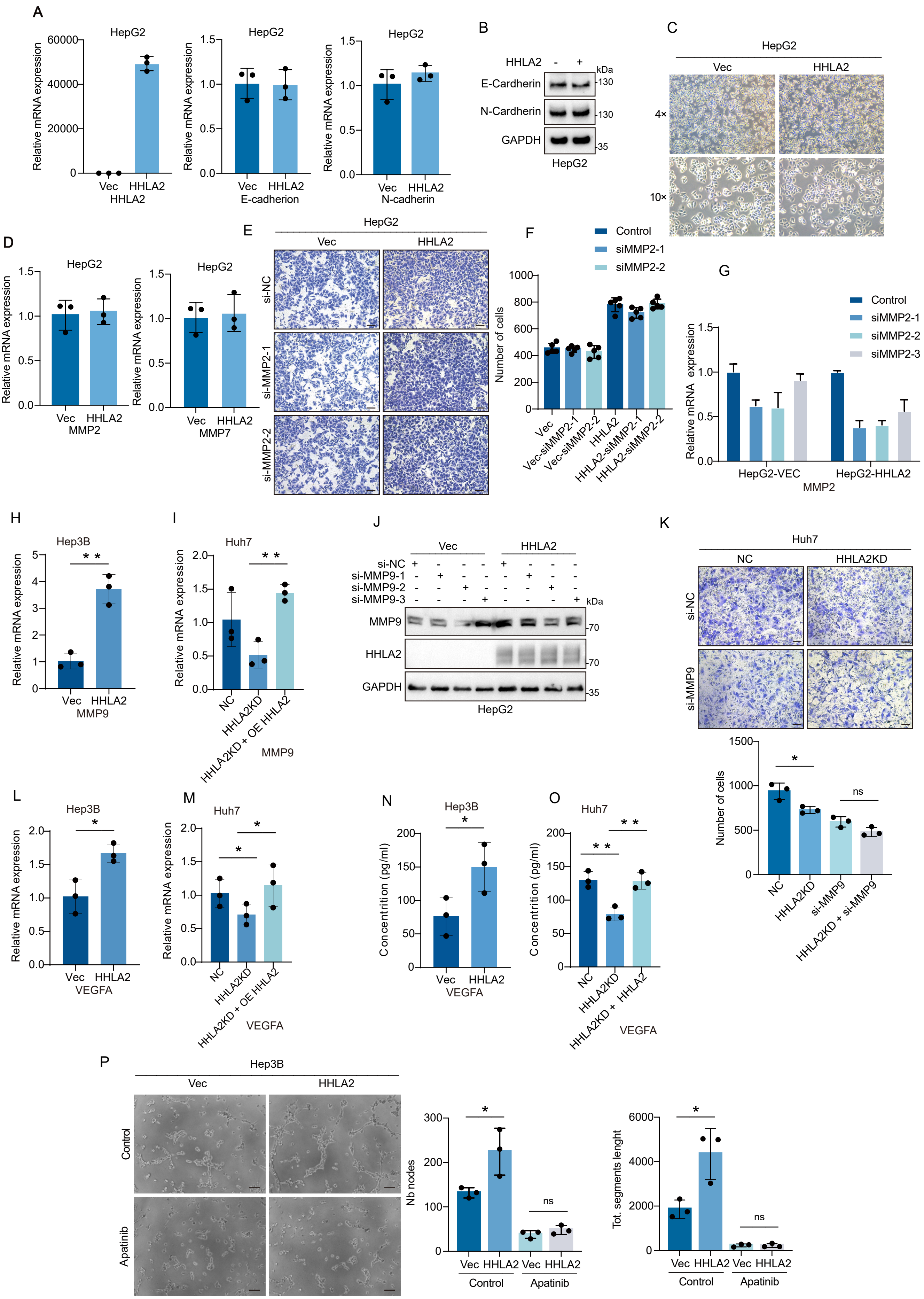
A
HepG2
Relative mRNA expression
Vec
HHLA2
E-cadherion
N-cadherin
B
HHLA2
E-Cardherin
N-Cardherin
GAPDH
kDa
HepG2
C
4×
10×
D
MMP2
MMP7
E
si-NC
si-MMP2-1
si-MMP2-2
F
Control
siMMP2-1
siMMP2-2
Number of cells
Vec-siMMP2-1
Vec-siMMP2-2
HHLA2-siMMP2-1
HHLA2-siMMP2-2
G
siMMP2-3
HepG2-VEC
HepG2-HHLA2
H
Hep3B
MMP9
I
Huh7
NC
HHLA2KD
HHLA2KD + OE HHLA2
J
si-NC
si-MMP9-1
si-MMP9-2
si-MMP9-3
MMP9
K
HHLA2KD
si-MMP9
HHLA2KD + si-MMP9
ns
L
VEGFA
M
N
Concentrition (pg/ml)
O
HHLA2KD + HHLA2
P
Control
Apatinib
Nb nodes
Tot. segments lenght

**Supplemental Figure 6. MMP family and VEGFA-VEGFR2 signaling contribute to HHLA2-driven invasion and tumor angiogenesis. (A–D)** qRT-PCR analysis of HHLA2, E-cadherin, and N-cadherin mRNA expression (A); Western blot analysis of the indicated protein levels (B); representative bright-field images showing morphology (C); and qRT-PCR analysis of MMP2 mRNA expression (D) in HepG2-Vec and HepG2-HHLA2 cells. **(E–G)** Representative images (E) and quantification (F) of invading HepG2-Vec and HepG2-HHLA2 cells with or without MMP2 knockdown; MMP2 knockdown efficacy was assessed by qRT-PCR (G). **(H, I)** qRT-PCR analysis of MMP9 mRNA expression in Hep3B-Vec and Hep3B-HHLA2 cells (H) and Huh7 cells with or without HHLA2 knockdown and rescue (I). **(J)** Western blot validation of MMP9 knockdown in HepG2-Vec and HepG2-HHLA2 cells. **(K)** Transwell invasion assays of Huh7 and Huh7-HHLA2KD cells with or without MMP9 knockdown. **(L, M)** qRT-PCR analysis of VEGFA mRNA expression in Hep3B cells with or without HHLA2 overexpression (L) and Huh7 cells with or without HHLA2 knockdown and rescue (M). **(N, O)** ELISA analysis of VEGFA secretion in Hep3B cells with or without HHLA2 overexpression (N) and Huh7 cells with or without HHLA2 knockdown and rescue (O). **(P)** HUVEC tube formation assays using conditioned media from Hep3B-Vec or Hep3B-HHLA2 cells, with or without treatment with 50 ng/ml apatinib (VEGF receptor inhibitor). P values were determined by two-tailed Student's t test. * $P < 0.05$, ** $P < 0.01$, *** $P < 0.001$, **** $P < 0.0001$. Scale bars, 100 μm.

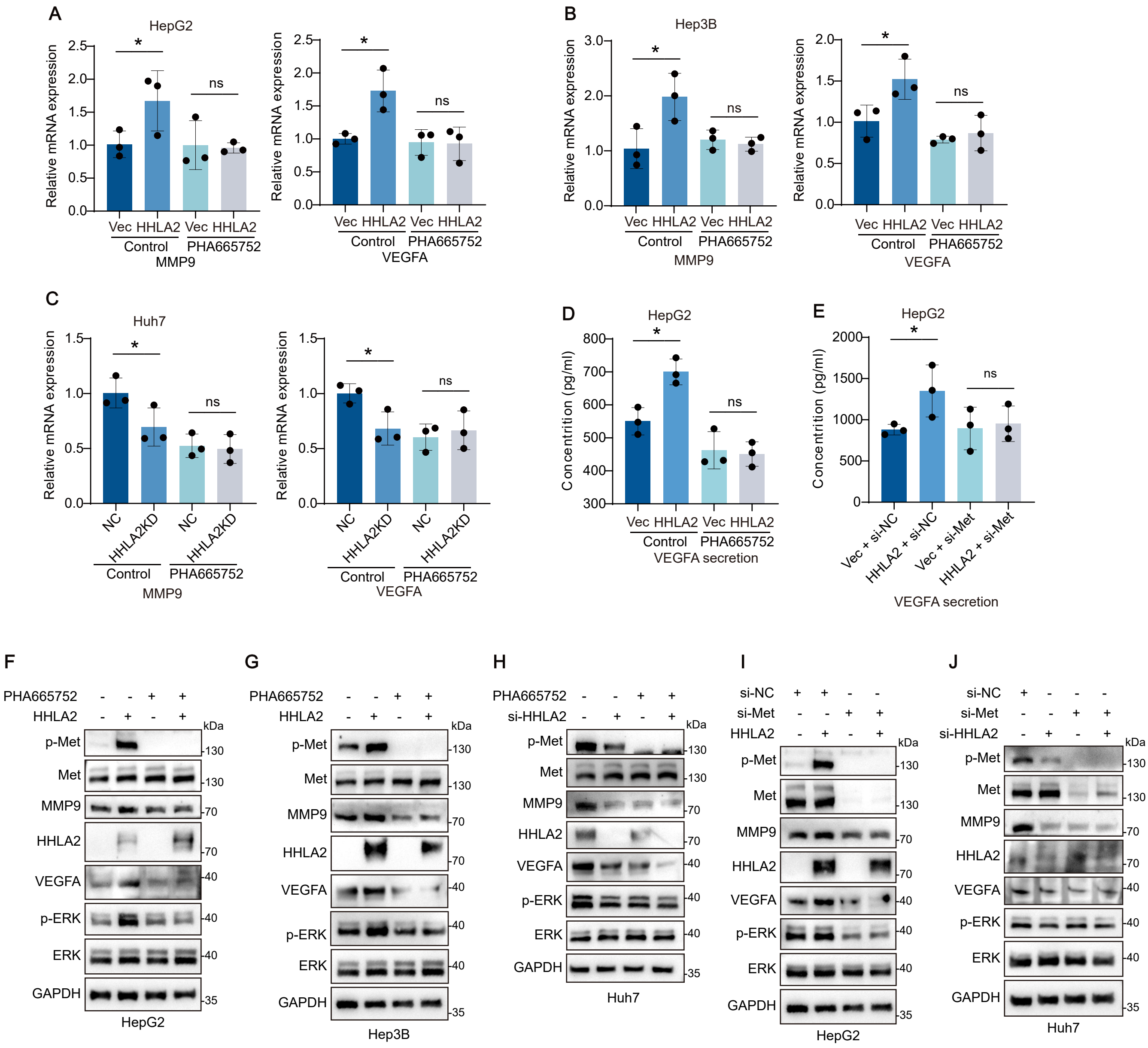


**Supplemental Figure 7. c-Met activation is required for HHLA2-mediated MMP9 and VEGFA expression and secretion. (A–C)** qRT-PCR analysis of MMP9 and VEGFA mRNA expression in (A) HepG2-Vec and HepG2-HHLA2 cells, (B) Hep3B-Vec and Hep3B-HHLA2 cells, and (C) Huh7-Vec and Huh7-HHLA2KD cells. All cell lines were treated with or without 100 nM PHA665752 (c-Met inhibitor). **(D, E)** ELISA of VEGFA secretion in HepG2-Vec and HepG2-HHLA2 cells with or without 100 nM PHA665752 treatment (D) and with or without c-MET knockdown (E). **(F–H)** Western blot analysis of ERK activation, VEGFA, and MMP9 expression in (F) HepG2-Vec and HepG2-HHLA2 cells, (G) Hep3B-Vec and Hep3B-HHLA2 cells, and (H) Huh7-Vec and Huh7-HHLA2KD cells. All cell lines were treated with or without 100 nM PHA665752. **(I, J)** Western blot analysis of ERK activation, VEGFA, and MMP9 expression in (I) HepG2-Vec and HepG2-HHLA2 cells and (J) Huh7-Vec and Huh7-HHLA2KD cells. All cell lines were treated with or without c-MET knockdown. P values were determined by two-tailed Student's t test. ns, not significant; * $P < 0.05$, ** $P < 0.01$, *** $P < 0.001$, **** $P < 0.0001$. Scale bars, 100 μm.

Fig. S8

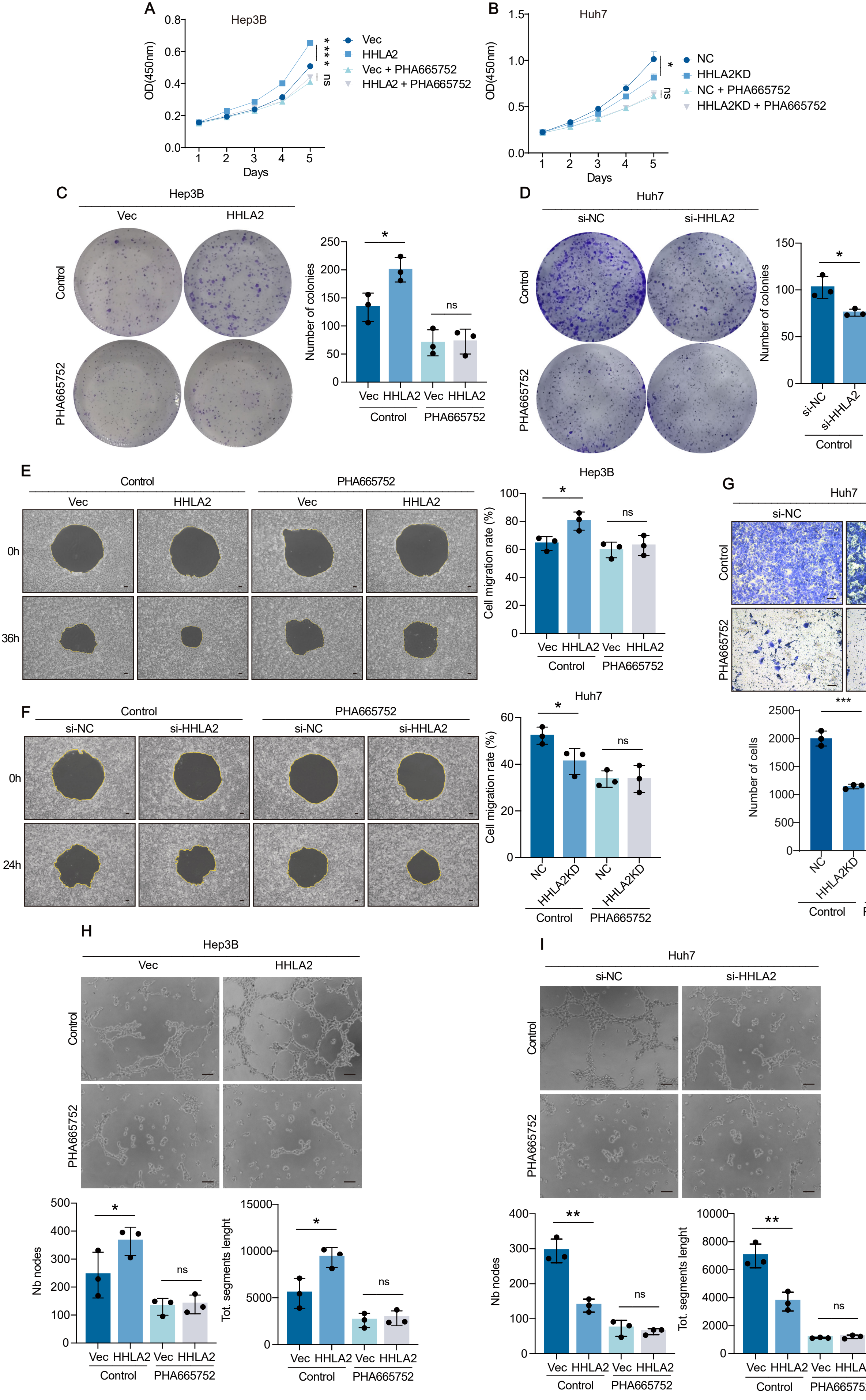
A
Hep3B
OD(450nm)
Days
Vec
HHLA2
Vec + PHA665752
HHLA2 + PHA665752
****
ns
B
Huh7
OD(450nm)
Days
NC
HHLA2KD
NC + PHA665752
HHLA2KD + PHA665752
*
ns
C
Hep3B
Vec
HHLA2
Control
PHA665752
Number of colonies
*
ns
Vec HHLA2 Vec HHLA2
Control
PHA665752
D
Huh7
si-NC
si-HHLA2
Control
PHA665752
Number of colonies
*
ns
si-NC si-HHLA2 si-NC si-HHLA2
Control
PHA665752
E
Control
PHA665752
Vec
HHLA2
Vec
HHLA2
0h
36h
Hep3B
Cell migration rate (%)
*
ns
Vec HHLA2 Vec HHLA2
Control
PHA665752
F
Control
PHA665752
si-NC
si-HHLA2
si-NC
si-HHLA2
0h
24h
Huh7
Cell migration rate (%)
*
ns
NC HHLA2KD NC HHLA2KD
Control
PHA665752
G
Huh7
si-NC
si-HHLA2
Control
PHA665752
Number of cells
***
ns
NC HHLA2KD NC HHLA2KD
Control
PHA665752
H
Hep3B
Vec
HHLA2
Control
PHA665752
Nb nodes
*
ns
Vec HHLA2 Vec HHLA2
Control
PHA665752
Tot. segments lenght
*
ns
Vec HHLA2 Vec HHLA2
Control
PHA665752
I
Huh7
si-NC
si-HHLA2
Control
PHA665752
Nb nodes
**
ns
Vec HHLA2 Vec HHLA2
Control
PHA665752
Tot. segments lenght
**
ns
Vec HHLA2 Vec HHLA2
Control
PHA665752

**Supplemental Figure 8. c-Met inhibition by PHA665752 abrogates HHLA2-mediated HCC progression in vitro. (A, B)** Cell proliferation assessed by CCK-8 assay in HepB3-Vec and HepB3-HHLA2 cells (A) or Huh7-Vec and Huh7-HHLA2KD cells (B). **(C, D)** Two-dimensional colony formation assay in HepB3-Vec and HepB3-HHLA2 cells (C) or Huh7-Vec and Huh7-HHLA2KD cells (D). **(E, F)** Cell migration assay in HepB3-Vec and HepB3-HHLA2 cells (E) or Huh7-Vec and Huh7-HHLA2KD cells (F). **(G)** Transwell invasion assay of Huh7-Vec and Huh7-HHLA2KD cells. **(H, I)** HUVEC tube formation assay induced by conditioned media from HepB3-Vec and HepB3-HHLA2 cells (H) or Huh7-Vec and Huh7-HHLA2KD cells (I). All cells were treated with 100 nM PHA665752 or DMSO as indicated. P values were determined by two-tailed Student's t test. ns, not significant; * P < 0.05, ** P < 0.01, *** P < 0.001, **** P < 0.0001. Scale bars, 100 μm.

Fig. S9

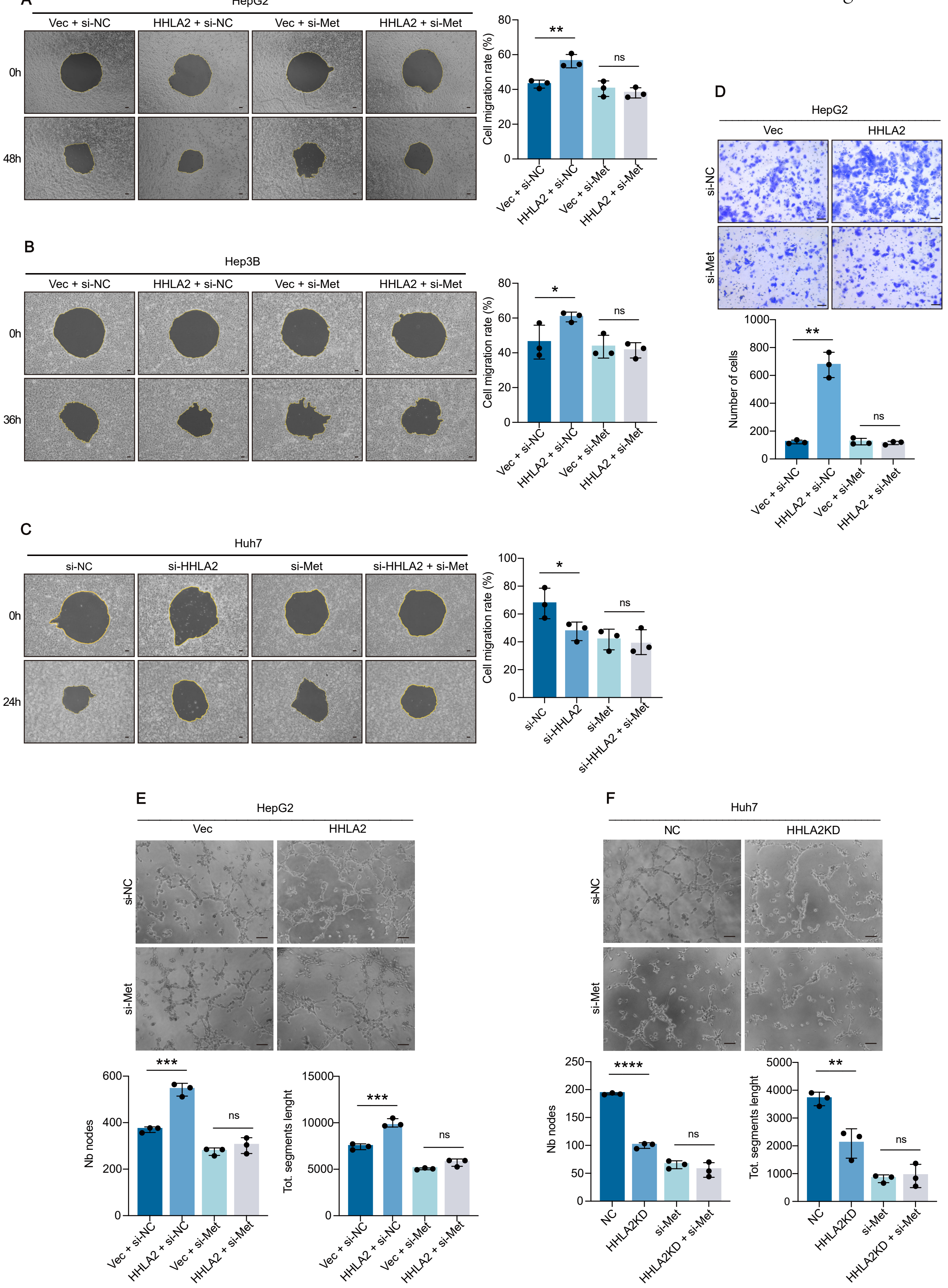
A
HepG2
Vec + si-NC
HHLA2 + si-NC
Vec + si-Met
HHLA2 + si-Met
0h
48h
Cell migration rate (%)
**
ns
B
Hep3B
Vec + si-NC
HHLA2 + si-NC
Vec + si-Met
HHLA2 + si-Met
0h
36h
Cell migration rate (%)
*
ns
C
Huh7
si-NC
si-HHLA2
si-Met
si-HHLA2 + si-Met
0h
24h
Cell migration rate (%)
*
ns
D
HepG2
Vec
HHLA2
si-NC
si-Met
Number of cells
**
ns
Vec + si-NC
HHLA2 + si-NC
Vec + si-Met
HHLA2 + si-Met
E
HepG2
Vec
HHLA2
si-NC
si-Met
Nb nodes
Tot. segments lenght
***
ns
F
Huh7
NC
HHLA2KD
si-NC
si-Met
Nb nodes
Tot. segments lenght
****
**
ns
NC
HHLA2KD
si-Met
HHLA2KD + si-Met

**Supplemental Figure 9. c-Met knockdown abrogates HHLA2-mediated HCC progression in vitro. (A–C)** Cell migration assays of HepG2-Vec and HepG2-HHLA2 cells (A), HepB3-Vec and HepB3-HHLA2 cells (B), or Huh7-Vec and Huh7-HHLA2KD cells (C) with or without c-MET knockdown. **(D)** Transwell invasion assay of HepG2-Vec and HepG2-HHLA2 cells with or without c-MET knockdown. **(E, F)** HUVEC tube formation assays induced by conditioned media from HepG2-Vec and HepG2-HHLA2 cells (E) or Huh7-Vec and Huh7-HHLA2KD cells (F) with or without c-MET knockdown. P values were determined by two-tailed Student's t test. ns, not significant; * $P < 0.05$, ** $P < 0.01$, **** $P < 0.0001$. Scale bars, 100 μm.

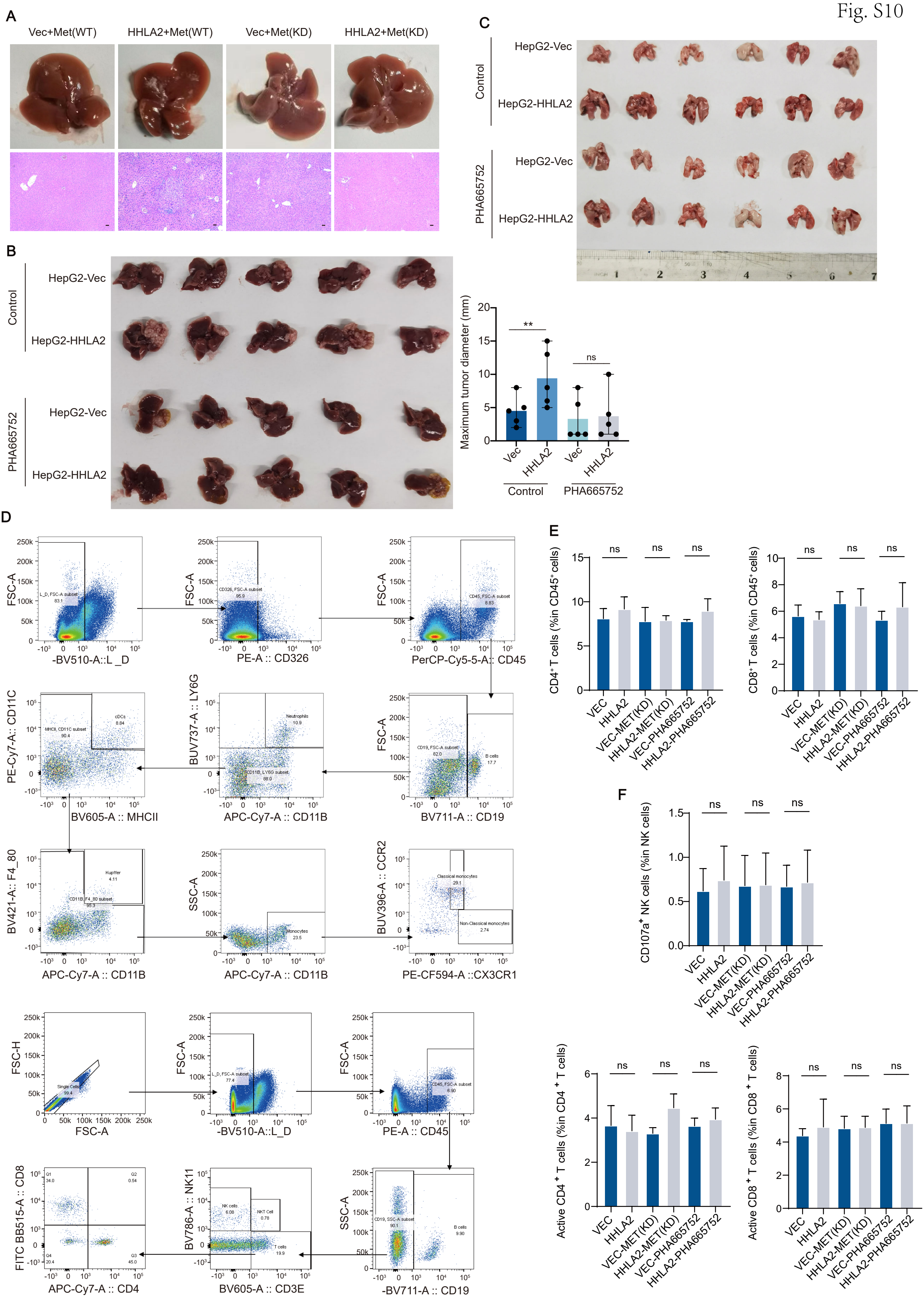
A
Vec+Met(WT)
HHLA2+Met(WT)
Vec+Met(KD)
HHLA2+Met(KD)
B
Control
PHA665752
HepG2-Vec
HepG2-HHLA2
Maximum tumor diameter (mm)
**
ns
Vec
HHLA2
C
D
-BV510-A::L_D
PE-A :: CD326
PerCP-Cy5-5-A:: CD45
PE-Cy7-A:: CD11C
BV605-A :: MHCII
BUV737-A :: LY6G
APC-Cy7-A :: CD11B
BV711-A :: CD19
BV421-A:: F4_80
SSC-A
BUV396-A :: CCR2
PE-CF594-A ::CX3CR1
FSC-H
FSC-A
PE-A :: CD45
FITC BB515-A :: CD8
APC-Cy7-A :: CD4
BV786-A :: NK11
BV605-A :: CD3E
-BV711-A :: CD19
E
CD4⁺ T cells (%in CD45⁺ cells)
CD8⁺ T cells (%in CD45⁺ cells)
F
CD107a⁺ NK cells (%in NK cells)
Active CD4⁺ T cells (%in CD4⁺ T cells)
Active CD8⁺ T cells (%in CD8⁺ T cells)
VEC
HHLA2
VEC-MET(KD)
HHLA2-MET(KD)
VEC-PHA665752
HHLA2-PHA665752
ns

**Supplemental Figure 10. Infiltration of immune cell subsets in mouse tumor tissue. (A)** Representative images of orthotopic liver tumors in C57BL/6 mice following HDTVi delivery of myc-HHLA2, a control vector, wild-type (WT) c-MET, or kinase-dead (KD) c-MET. **(B)** Representative images of liver tumors in orthotopic xenograft models in nude mice injected with HepG2-HHLA2 cells, followed by intraperitoneal administration of PHA665752 (n = 5 mice per group). **(C)** Representative images of lung tissues from nude mice following tail vein injection of HepG2-Vec or HepG2-HHLA2 cells, with or without PHA665752 treatment every other day for 28 days. Lung tissues were assessed for HCC metastasis. **(D–F)** Flow cytometry analysis of immune cell subpopulations in C57BL/6 mouse livers following HDTVi delivery of Myc-HHLA2 or a control vector, along with N-RasV12/myr-AKT1 and the Sleeping Beauty transposon system. (D) Representative flow cytometry plots showcasing various immune cell populations (macrophages, neutrophils, CD4+ T cells, CD8+ T cells, and B cells) in the liver. (E) Quantification of the relative abundance of each immune cell subpopulation. (F) Changes in indicated immune cell activity in different treatment groups. Data are presented as mean ± SD (n = 5 mice per group). P values were determined by two-tailed Student’s t test. ns, not significant; * $P < 0.05$, ** $P < 0.01$, **** $P < 0.0001$. Scale bars, 100 μm.

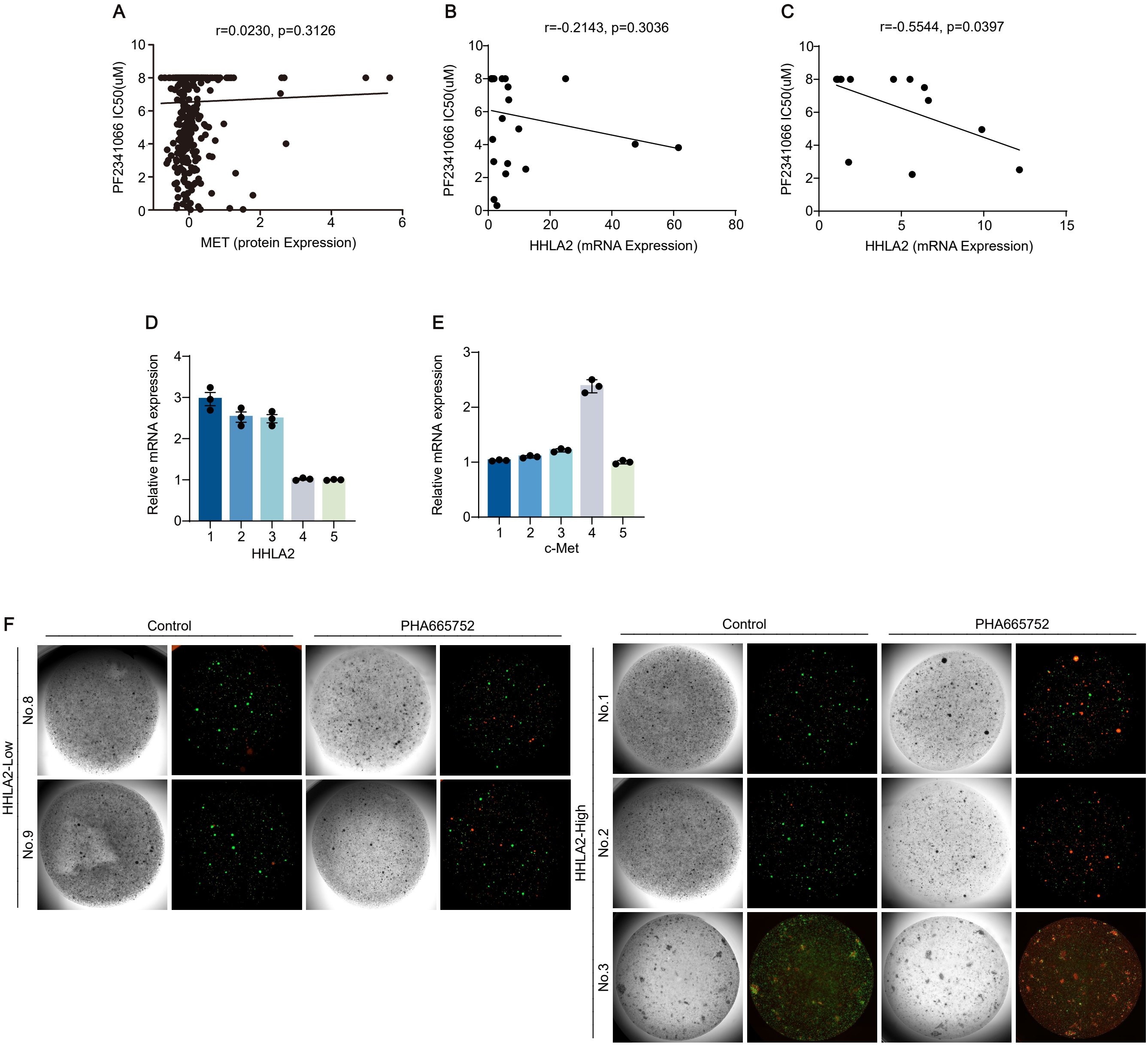


**Supplemental Figure 11. HHLA2 expression predicts efficacy of c-Met inhibitor therapy in HCC. (A–C)** Correlation between c-Met/HHLA2 expression and c-Met inhibitor efficacy in HCC cell lines. Relationship between c-Met expression level (A), HHLA2 expression level (B), or HHLA2 expression levels with abnormal c-Met expression (C) and the efficacy of the c-Met inhibitor PF2341066 in cancer cell lines from the Cancer Cell Line Encyclopedia (CCLE) database. **(D, E)** Relative HHLA2 (D) and c-Met (E) mRNA expression in five HCC organoids determined by qRT-PCR. **(F)** Representative images showing patient-derived organoid (PDO) death in response to c-Met inhibitor PHA665752 treatment. Dead cells are stained red with propidium iodide (PI), and live cells are stained green with Calcein-AM.
